# Rationally designed multimeric nanovaccines using icosahedral DNA origami for molecularly controlled display of SARS-CoV-2 receptor binding domain

**DOI:** 10.1101/2023.08.24.554561

**Authors:** Qingqing Feng, Keman Cheng, Lizhuo Zhang, Xiaoyu Gao, Jie Liang, Guangna Liu, Nana Ma, Chen Xu, Ming Tang, Liting Chen, Xinwei Wang, Xuehui Ma, Jiajia Zou, Quanwei Shi, Pei Du, Qihui Wang, Guangjun Nie, Xiao Zhao

## Abstract

Multivalent antigen display on nanoparticles can enhance the immunogenicity of nanovaccines targeting viral moieties, such as the receptor binding domain (RBD) of SARS-CoV-2. However, particle morphology and size of current nanovaccines are significantly different from those of SARS-CoV-2. Additionally, surface antigen patterns are not controllable to enable the optimization of B cell activation. Herein, we employed an icosahedral DNA origami (ICO) as a display particle for SARS-CoV-2 RBD nanovaccines. The morphology and diameter of the particles were close to those of the virus (91 ± 11 nm). The surface addressability of the DNA origami permitted facile modification of the ICO surface with numerous RBD antigen clusters (ICO-RBD) to form various antigen patterns. Using an *in vitro* screening system, we demonstrate that the antigen spacing, antigen copies within clusters and cluster number parameters of the surface antigen pattern all impact the ability of the nanovaccines to activate B cells. Importantly, the optimized ICO-RBD nanovaccines evoked stronger and more enduring humoral and T cell immune responses in mouse models compared to soluble RBD antigens. Our vaccines activated similar humoral immunity and slightly stronger cellular immunity compared to mRNA vaccines. These results provide reference principles for the rational design of nanovaccines and exemplify the utility of DNA origami as a display platform for vaccines against infectious disease.

## Introduction

The global coronavirus pandemic caused by the severe acute respiratory syndrome coronavirus 2 (SARS-CoV-2), has profoundly affected human health^1^. Several types of vaccines fabricated via differing technological routes were developed to combat the coronavirus disease 2019 (COVID-19) pandemic caused by SARS-CoV-2^2, 3^. Among them, the subunit vaccines that comprise only key viral proteins have become the most widely successful SARS-CoV-2 vaccine candidates, in both clinical and preclinical stages, due to their well-characterized ingredients and favorable safety^4–6^. The receptor binding domain (RBD) in the surface spike (S) protein of SARS-CoV-2 is the dominant domain for the interaction between viruses and human cells, which makes this subunit the most attractive vaccine epitope target because it elicits functionally neutralizing antibodies to block virus invasion, avoiding the potential risk of antibody-dependent enhancement due to non-neutralizing antibodies^7–9^.

To date, several subunit vaccines targeting SARS-CoV-2 RBD have been approved or entered clinical trials. Although there have been seminal reports about RBD monomer vaccines^10, 11^, the low molecular weight and surface valence limit their immunogenicity. RBD multimerization, such as dimerization via Fc fragment fusion or disulfide bonding and trimerization via stabilization by the T4 fibritin foldon domain, is a popular and useful strategy to overcome these drawbacks^12–16^. Recently, to further improve antigen size and surface valence, the method of displaying multiple copies of RBD on nanoparticles has received increasing attention. Nanoparticles with highly repetitive surface patterns, such as virus-like particles and ferritin, have been employed to establish multimeric RBD nanovaccines to improve the lymph node drainage of antigens and promote the ability to induce B cell receptor (BCR) crosslinking and activation^17–19^. However, the sizes of these nanovaccines (10-30 nm) are significantly smaller than that of SARS-CoV-2 (∼90 nm), and the surface antigen patterns differ from the cluster distribution in SARS-CoV-2^20–23^. Nanoparticles with controllable dimensional morphologies and surface patterns are required to improve the similarity between RBD nanovaccines and SARS-CoV-2 and to explore the structure-activity relationship between surface antigen pattern and vaccine efficacy.

As a type of DNA nanotechnology, DNA origami entails the combination of one or more template strands with numerous staple strands, which together self-assemble via complementary base pairing^24^. Due to the precise controllability of DNA strands, DNA origami is morphologically programmable and can be assembled into nanostructures with different sizes and morphologies^25, 26^. In addition, DNA origami possesses surface addressability because of the specificity of DNA sequences; the staple strands in DNA origami can be designed as capture strands, through sequence elongation and end modification, for subsequent molecular linkage with controllable spacing, number and pattern^25^. Due to these characteristics, DNA origami has become an ideal tool to study the relationship between the spatial arrangement and biological function of proteins, such as enzymes, receptors/ligands and antibodies, at the nanoscale^27–30^. Importantly, nanoscale antigen organization has been proven to affect B cell activation^31^. Additionally, the surface pattern of RBD distribution is associated with SARS-CoV-2 infection ability^32^. Therefore, using DNA origami to build biomimetic nanovaccines with similar sizes and surface antigen patterns to the corresponding viruses, such as SARS-CoV-2, may facilitate the development of highly effective antiviral vaccines.

Although there were several studies about the SARS-CoV-2 nanovaccines based on the DNA origami scaffolds, the structure-activity relationship between surface antigen pattern and vaccine efficacy was still elusive, and the underlying mechanism of those immune effects *in vivo* has not been scrutinized^33, 34^. In the present study, we employed DNA origami to rationally design and precisely assemble RBD nanovaccines against SARS-CoV-2. We first constructed an icosahedral DNA origami (ICO) of ∼90 nm in diameter, which approximated the morphology and size of SARS-CoV-2 (**Figure 1a**). We then generated various surface antigen patterns with different parameters, including antigen spacing, antigen copies within clusters and cluster numbers, and precisely decorated RBD antigens onto ICO (ICO-RBD) using an “engraving-printing” strategy (**Figure 1a**). *In vitro* experiments using B cells stably expressing the RBD antigen’s cognate IgG receptor were performed to interrogate the impacts of these parameters on B cell activation and screen for nanovaccines with optimal surface RBD patterns (**Figure 1b**). Finally, we systematically evaluated the short- and long-term immune effects of the optimized ICO-RBD nanovaccines in mouse models and compared the immune effects of the nanovaccines with mRNA vaccines (**Figure 1b**).

**Figure 1.**
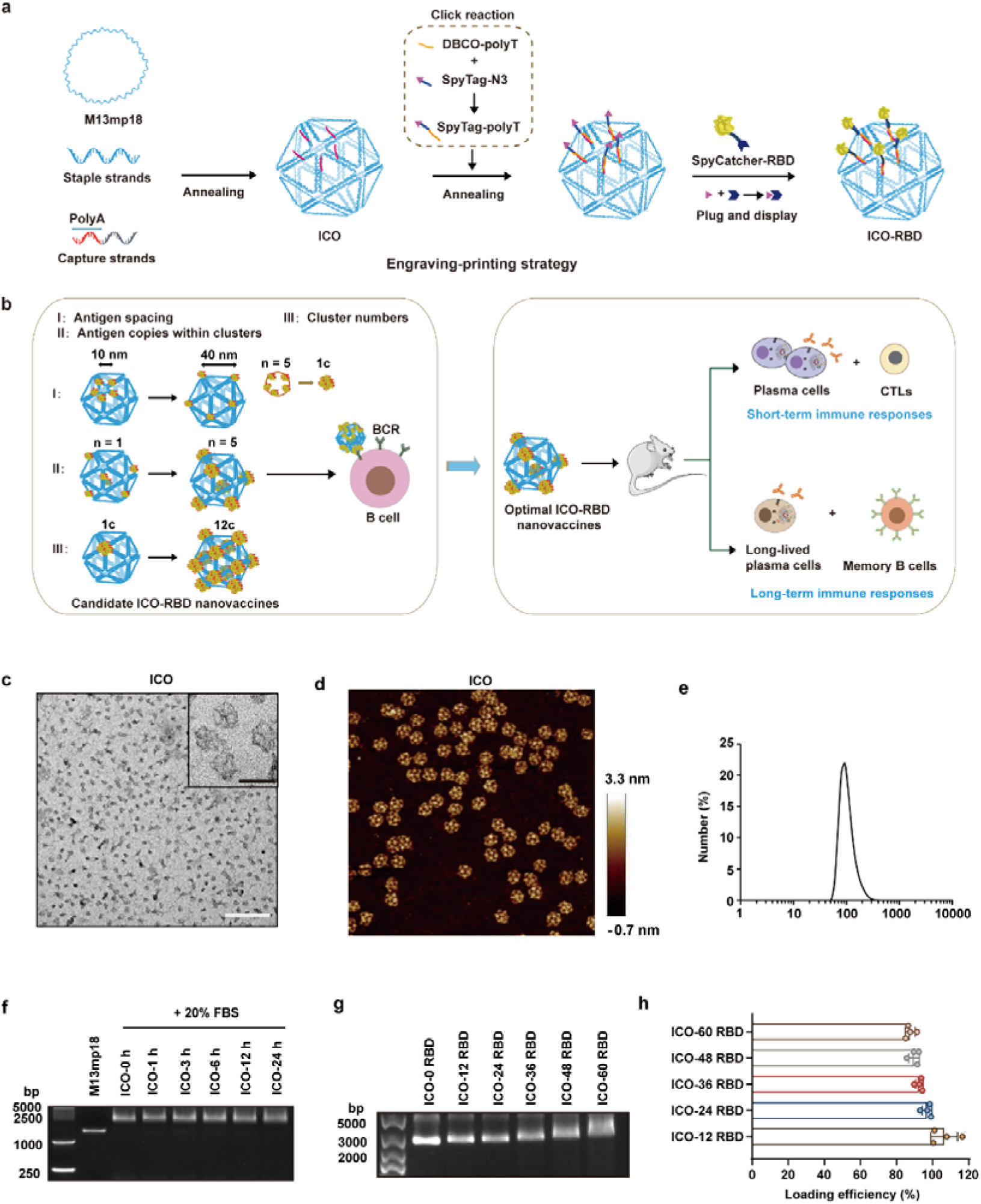
Design and construction of ICO-RBD nanovaccines. **(a)** Schematic illustration of the “engraving-printing” strategy to decorate RBD antigens onto ICO. The genomic DNA of M13mp18 bacteriophage was annealed and folded together with staple and capture strands (with polyA overhangs) to form the ICO nanocore. The SpyTag-N3 and DBCO-polyT were covalently linked through click chemistry to form the SpyTag-polyT conjugates. Next, the SpyTag peptides were engraved onto ICO through hybridization between polyT and polyA. The fusion proteins, SpyCatcher-RBD (prototype), were printed onto ICO through the spontaneous covalent isopeptide bond formation between SpyCatcher and SpyTag. **(b)** Schematic illustration of the optimization screening of surface antigen patterns based on the efficiency of the nanovaccine to activate RBD-specific IgG-BCR *in vitro* and the evaluation of efficient short- and long-term immune responses *in vivo*. Various surface antigen patterns with different parameters, including antigen spacing (10–40 nm), antigen copies within clusters (1-5) and cluster numbers (1-12), were screened. **(c-e)** Characterization of ICO, including TEM (c), AFM (d) and DLS (e) analysis. Scale bar (white), 1 μm. Scale bar (black), 100 nm. **(f)** Gel electrophoresis of genomic DNA of M13mp18 bacteriophage and ICO after incubation in 20% FBS for different time. **(g-h)** Gel electrophoresis (g) and loading efficiency of ICO-RBD nanovaccines surface-modified with 12, 24, 36, 48 or 60 RBD antigens (n = 4) (h). The data were processed on GraphPad Prism 8 and are presented as the mean ± SD.

## Results

### Design and construction of ICO-RBD nanovaccines

The ICO was assembled by slowly annealing the genomic DNA of M13mp18 bacteriophage as a template strand together with multiple staple and capture strands (**Supplementary Data 1**). In negative staining transmission electron microscopy (TEM) and atomic force microscopy (AFM), ICO exhibited a uniform, hollowed-out polyhedron structure of ∼80–90 nm in diameter (**Figures 1c-d**). The hydrophilic diameter of ICO was 93 nm, as detected using dynamic light scattering (DLS; **Figure 1e**), which is close to that of SARS-CoV-2. We also verified the successful assembly of ICO according to migration in agarose gel electrophoresis, compared to the genomic DNA of M13mp18 bacteriophage (**Figure 1f**). In addition, ICO exhibited high stability property under 20% fetal bovine serum (FBS) up to 24 h (**Figure 1f**), which is favorable to stimulate immune cells *in vivo*.

Next, we adopted an “engraving-printing” strategy to decorate prototype RBD antigens onto ICO (**Figure 1a**). In brief, the 5’-ends of staple strands at predetermined positions were elongated with overhanging polyA to form capture strands. The polyA overhang did not interfere with origami assembly, even if the number of capture strands increased to 60 (**Supplementary Figure 1a**). Next, the C-terminal azide-modified SpyTag peptide glue (SpyTag-N3) and 5’-end dibenzocyclooctyne (DBCO)-modified polyT (DBCO-polyT) were covalently linked through click chemistry to form SpyTag-polyT conjugates. Next, we “engraved” ICO with the SpyTag peptides through DNA complementary hybridization between the polyT in the SpyTag-polyT conjugates and the polyA overhang in the capture strands (**Supplementary Figure 1b**). Finally, fusion proteins of SpyCatcher and RBD (N-SpyCatcher-RBD-C; **y8**) were “printed” onto ICO through the spontaneous covalent isopeptide bond formation between SpyCatcher and SpyTag^35^, ultimately achieving site-specific decoration of RBD antigens onto the surface of ICO. We designed and constructed 5 different ICO-RBD nanovaccines surface-modified with 12, 24, 36, 48 or 60 RBD antigens. As the number of modified RBD antigens increased, the electrophoretic migration of ICO-RBD decreased (**Figure 1g**), indicating successful protein decoration of the nanoparticles. By measuring the DNA and protein concentration, we show that the loading efficiency of RBD antigens in ICO-RBD reached greater than 85% when the number of RBD antigens increased from 12 to 60 (**Figure 1h**). This high efficiency is attributed to the mild conditions and rapid reaction kinetics of the click chemistry reaction and peptide glue technology. In addition, the site-specific conjugation based on the peptide glue technology ensured a uniform outward orientation of RBD antigens on the ICO-RBD nanovaccines, which is beneficial to the antigenic immune stimulation of B cells.

### Differential efficacy of ICO-RBD nanovaccines with different surface antigen patterns on B cell activation

Next, we investigated the relationship between nanovaccine surface antigen pattern and B cell activation efficiency. We designed the surface RBD patterns in ICO-RBD nanovaccines in consideration of the cluster distribution features of RBD antigens in SARS-CoV-2. The precise control afforded by the DNA origami scaffold enabled us to fabricate ICO-RBD nanovaccines with varying antigen spacing, antigen copy number within clusters, and cluster numbers. We termed the different ICO-RBD nanovaccines ICO-RBD-xc×n-d nm, in which xc represents cluster number, n represents antigen copy number within clusters and d represents the antigen spacing within clusters. To assess the capacity of the nanovaccines to activate B cells, we generated B cells that express BCRs with the RBD-targeting monoclonal neutralizing antibody COVA2-15 (B-RBD cells), as previously described^36, 37^, and assessed the cell binding and Ca^2+^ influx induced by the different ICO-RBD nanovaccines *in vitro*.

We began with ICO-RBD nanovaccines with only 1 RBD cluster of 5 RBD copies within the cluster, but with different RBD spacing from 10 to 40 nm (**Figure 2a and Supplementary Figures 3a-b**). With decreased RBD spacing, the affinity between the ICO-RBD nanovaccines and B-RBD cells gradually increased (**Figure 2b**). The ability of ICO-RBD nanovaccines to trigger Ca^2+^ influx of B-RBD cells also negatively correlated with the RBD spacing (**Figure 2c**). When the RBD spacing reached 40 nm, the ability of ICO-RBD nanovaccines to activate Ca^2+^ influx of B-RBD cells was similar with that of soluble RBD monomers (**Figure 2c**). These results indicate that antigen clustering indeed enhances B cell activation over soluble RBD monomers, while the strongest B cell activation occurred under close antigen spacing. When the antigen spacing increases past a critical threshold, the RBD antigens lose their clustered characteristic.

**Figure 2.**
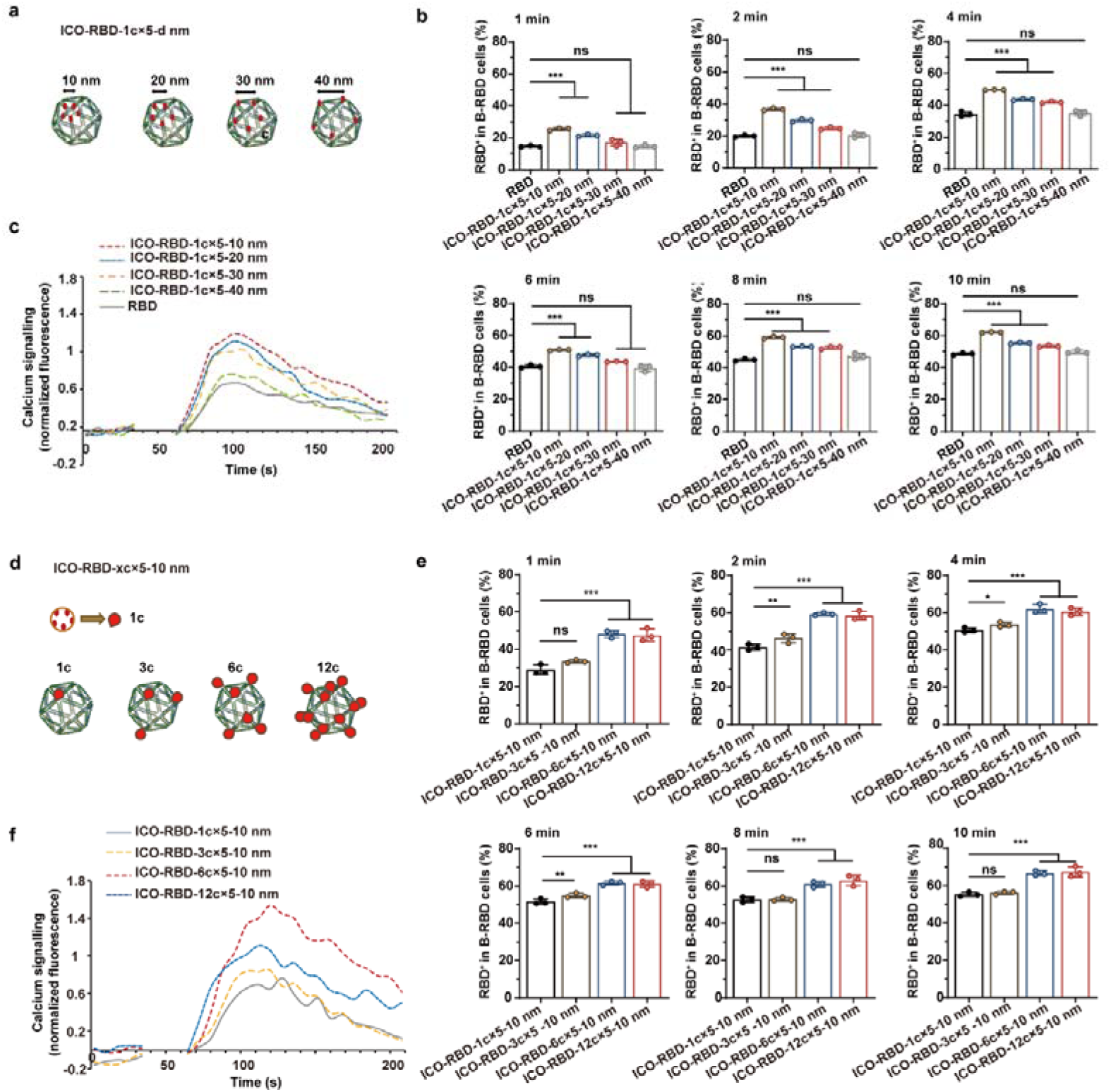
Effects of antigen spacing and cluster number on the efficiency of ICO-RBD nanovaccine activation of B-RBD cells. **(a)** Design of ICO-RBD nanovaccines with 1 RBD cluster of 5 RBD copies within clusters, but different RBD spacing from 10 to 40 nm. **(b)** Binding affinity between the SpyCatcher-RBD proteins (RBD) or ICO-RBD nanovaccines with different RBD spacing and B-RBD cells (n = 3). ICO-RBD nanovaccines bound to B-RBD cells were detected using anti-His antibodies and flow cytometry. **(c)** Ca^2+^ traces in B-RBD cells triggered by SpyCatcher-RBD proteins (RBD) or ICO-RBD nanovaccines with differing RBD spacing, as detected by Fluo-4 AM labeling and flow cytometry. **(d)** Design and AFM images of ICO-RBD nanovaccines with 1–12 RBD clusters of 5 RBD copies within clusters and 10 nm RBD spacing. **(e)** Binding affinity between the ICO-RBD nanovaccines with different RBD cluster numbers and B-RBD cells (n = 3). ICO-RBD nanovaccines bound to B-RBD cells were detected using anti-His antibodies and flow cytometry. **(f)** Ca^2+^ traces in B-RBD cells triggered by ICO-RBD nanovaccines with the different RBD cluster numbers, as detected by Fluo-4 AM labeling and flow cytometry measurement. The data were processed on GraphPad Prism 8 and are presented as the mean ± SD. Statistical significance (*P* value) was calculated by one-way ANOVA followed by Tukey’s test. *, *P* < 0.05; **, *P* < 0.01; ***, *P* < 0.001. ns, *P* > 0.05, no significant difference.

We next sought to define the impact of antigen cluster number in ICO-RBD nanovaccines on B cell activation. To this end, we fixed the number of antigen copies within clusters at 5 and the antigen spacing at 10 nm, while altering the cluster number from 1 to 12 (**Figure 2d and Supplementary Figures 3c-d**). As shown in **Figure 2e**, increasing the antigen cluster number improved the binding capacity between ICO-RBD nanovaccines and B-RBD cells. When the antigen cluster number reached 6, the binding ability became saturated, as indicated by the similar B cell binding of ICO-RBD-6c×5-10 nm and ICO-RBD-12c×5-10 nm. In the ICO-RBD nanovaccines with different antigen cluster numbers, ICO-RBD-6c×5-10 nm was also the most effective at activating Ca^2+^ influx in B-RBD cells (**Figure 2f**). This observation is also supported by the literature^23^, in which it has been reported that a surface multi-cluster pattern similar to that on the surface of SARS-CoV-2 is beneficial for immune stimulation.

To further investigate the role of antigen copy number within the clusters on B cell response, we programmed several ICO-RBD-6c×n-10 nm nanovaccines with 1 to 5 antigen copies within clusters (**Supplementary Figures 3e-f and 4a**). As shown in **Supplementary Figures 4b-c**, the cluster antigen copy number was also an important factor in the immune stimulation ability of ICO-RBD nanovaccines. Interestingly, although it possessed the same number of antigens as SARS-CoV-2, the ICO-RBD-6c×3-10 nm nanovaccine with 3 antigen copies within clusters was not the most effective formulation; nanovaccine efficacy continued to improve as cluster antigen copy number increased, demonstrating the importance of rational artificial vaccine design. In summary, through screening and functional evaluation, the ICO-RBD-6c×5-10 nm nanovaccine was the optimal ICO-RBD preparation. We chose these parameters for the subsequent *in vivo* experiments.

### Robust humoral immune responses induced by ICO-RBD nanovaccines *in vivo*

To assess the immunogenicity of ICO-RBD nanovaccines *in vivo*, we immunized 6-week-old BALB/c mice with two intramuscular injections of different ICO-RBD nanovaccines, containing 1 μg equivalent prototype RBD proteins, formulated with AddaVax adjuvant^6^ (**Figure 3a**). Phosphate buffered saline (PBS), RBD monomers (SpyCatcher-RBD proteins), ICO and RBD + ICO (physical mixture) were used as controls. As evaluated by enzyme-linked immunosorbent assay (ELISA), the serum titers of SARS-CoV-2 RBD-specific immunoglobulin G (IgG) in the RBD and ICO-RBD-6c×5-10 nm groups reached 148 and 1738, respectively, 2 weeks after the priming immunization (**Figure 3b**). After the boost immunization (given at week 3), the RBD-specific IgG titers elicited by ICO-RBD-6c×5-10 nm were greater than 1 × 10^5^, while the RBD monomers, RBD + ICO and other ICO-RBD nanovaccines, including ICO-RBD-1c×5-10 nm, ICO-RBD-1c×5-40 nm and ICO-RBD-6c×1-10 nm, produced only 1 × 10^3^-10^4^ RBD-specific IgG titers (**Figure 3c**). We monitored the RBD-specific IgG titers for 14 weeks; the time-titer curves in **Figure 3d** show that the RBD-specific IgG titers in the ICO-RBD-6c×5-10 nm group peaked at week 8 and only decreased slightly at week 14. However, the RBD-specific IgG titers in the other groups peaked at week 5 and decreased dramatically by week 14 (**Figure 3d**). We used the sera collected at week 5 and 14 in pseudovirus neutralization testing to analyze the neutralizing antibodies, as represented by the inhibitory dilution value that achieved 50% neutralization (ID_50_). Consistent with the RBD-specific IgG titer results, ICO-RBD-6c×5-10 nm had ID_50_ values over 100 times greater than those in the free RBD group (**Figures 3e-f**). Taken together, our data demonstrate that the ICO-RBD-6c×5-10 nm nanovaccines were highly immunogenic compared to RBD monomers.

**Figure 3.**
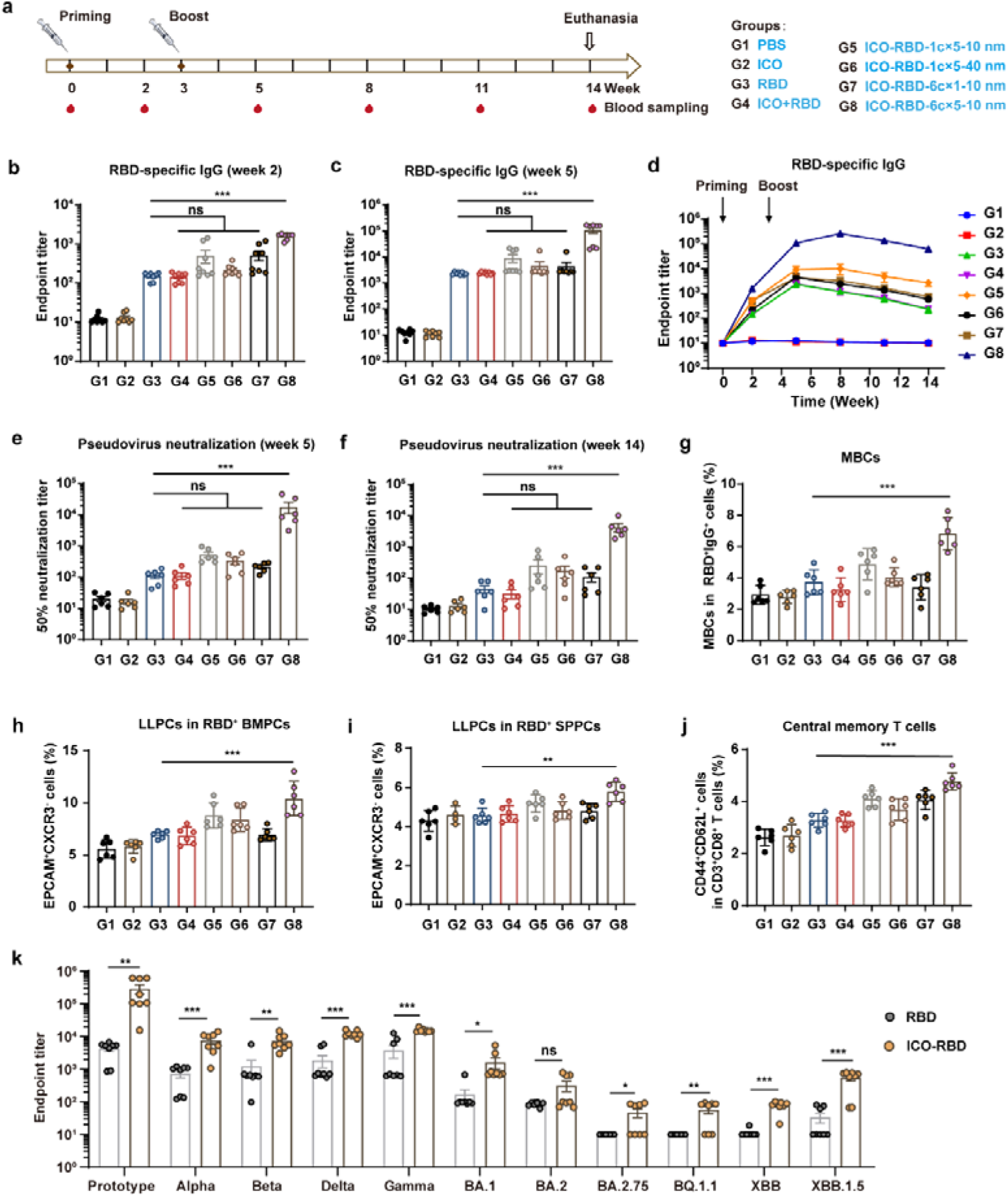
Humoral immune responses in ICO-RBD nanovaccine-immunized BALB/c mice. **(a)** Schematic illustration of the vaccination schedule and grouping information. The mice were randomly divided into 8 groups and were priming- and boost-vaccinated with the indicated vaccines containing prototype RBD at week 0 and 3, respectively. The red drop symbols indicate blood sampling and sera collection. All mice were euthanized at week 14 at which time memory immune cells were evaluated. **(b-c)** SARS-CoV-2 RBD-specific IgG titers in serum, measured at week 2 (b) and 5 (c) by ELISA (n = 8). Spycatcher-RBD proteins (prototype) were used to coat 96-well ELISA plates. **(d)** Time-titer curves of RBD-specific IgG titers measured every 3 weeks (n = 8). **(e-f)** Pseudovirus neutralization test of the sera collected at week 5 and 14, as represented by the inhibitory dilution value to achieve 50% neutralization (ID_50_; n = 6). **(g)** Percentages of MBCs (B220^+^CD38^+^) in RBD^+^IgG^+^ cells in splenocytes at week 14, as detected by flow cytometry (n = 6). **(h-i)** Percentages of LLPCs (EpCAM^hi^CXCR3^−^) in RBD^+^ bone marrow plasma cells (BMPCs, B220^-^CD138^+^) and RBD^+^ splenic plasma cells (SPPCs, B220^-^CD138^+^) at week 14, as detected using flow cytometry (n = 6). **(j)** Percentages of central memory T cells (CD44^+^CD62L^+^) in CD3^+^CD8^+^ T cells in splenocytes at week 14, as detected using flow cytometry (n = 6). **(k)** Serum titers of IgG against variants, measured at week 5 by ELISA (n = 8). Different RBD proteins were used to coat 96-well ELISA plates, including prototype (Wuhan-Hu-1 strain), Alpha [B.1.1.7], Beta [B.1.351], Gamma [P.1], Delta [B.1.617.2] and Omicron [BA.1, BA.2, BA.2.75, BQ.1.1, XBB and XBB.1.5]. The data are presented as the mean ± SEM in panels b-f and k, and the mean ± SD in panels g-j. The data were processed on GraphPad Prism 8. Statistical significance (*P* value) was calculated by one-way ANOVA followed by Tukey’s test. **, *P* < 0.01; ***, *P* < 0.001. ns, *P* > 0.05, no significant difference.

Next, to assess whether the ICO-RBD nanovaccines can induce long-lasting immune effects, we evaluated the memory immune cells, including memory B cells (MBCs), long-lived plasma cells (LLPCs), effector memory T cells and central memory T cells, at week 14^38, 39^. Compared with RBD monomers, the ICO-RBD-6c×5-10 nm nanovaccines induced a significantly greater number of RBD-specific IgG^+^ MBCs (**Figure 3g**). The proportions of LLPCs in the RBD-specific bone marrow plasma cells and splenic plasma cells were also greater in the ICO-RBD-6c×5-10 nm group than in the RBD group (**Figures 3h-i and Supplementary Figures 5a-b**). In addition, compared with mice in the RBD group, although effector memory T cells were not significantly induced (**Supplementary Figure 5c**), central memory T cells were significantly increased in the splenocytes from ICO-RBD-6c×5-10 nm nanovaccine-immunized mice (**Figure 3j**).

Finally, to assess the broad spectrum of RBD monomers and ICO-RBD-6c×5-10 nm nanovaccines containing prototype RBD, we used a panel of SARS-CoV-2 RBD antigens to test the serum titers of IgG against the protype and the variants, including prototype (Wuhan-Hu-1 strain), Alpha [B.1.1.7], Beta [B.1.351], Gamma [P.1], Delta [B.1.617.2] and Omicron [BA.1, BA.2, BA.2.75, BQ.1.1, XBB and XBB.1.5]. As shown in **Figure 3k**, the ICO-RBD-6c×5-10 nm nanovaccines induced more IgG against all variants than the RBD monomers. In particular, for Omicron [BA.1], [BA.2] and [XBB.1.5], the IgG titers in the ICO-RBD-6c×5-10 nm nanovaccine group reach to 1608, 315 and 544, respectively, being significantly higher than 168, 86 and 33 in the RBD group (**Figure 3k**). This is an interesting phenomenon, since the immune escape against Omicron variants is a widespread problem in the prototype RBD vaccines. By optimized assembly on the DNA origami scaffold, the ICO-RBD nanovaccine (prototype) can activate an enhanced antibody response against Omicron variants, which is important for the prevention of SARS-CoV-2 as the variants continue to emerge.

In summary, the DNA origami-based nanovaccines induced markedly increased neutralization antibodies against RBD and 1.5-fold more RBD-specific memory immune cells than RBD monomers, which is expected to provide a persistent protective humoral response.

### *In vivo* induction of T cell immune responses by ICO-RBD nanovaccines

There are numerous CD4^+^ and CD8^+^ T cell epitopes in the RBD, which are also critical for virus defense and clearance^40, 41^. Next, to measure the T cell responses induced by the ICO-RBD nanovaccine (ICO-RBD-6c×5-10 nm), we performed intracellular cytokine staining assays to analyze the different T cell populations within the spleen and lung at week 3 and 6, respectively. As shown in **Figures 4a-d**, the ICO-RBD nanovaccine elicited more RBD-specific CD8^+^ and CD4^+^ T cells expressing interferon-γ (IFN-γ), interleukin (IL)-2 and tumor necrosis factor-α (TNF-α) than RBD monomers, in both spleen and lung tissue, indicating that ICO-RBD nanovaccines induced strong type 1 T helper cell (Th1)-biased immune responses *in vivo*. IL-4 expressing CD4^+^ T cells were rarely induced across the three groups after priming and boost immunization, indicating that Th2-biased immune cells were not evoked (**Figures 4e-f**). Tissue resident memory T cells (TRMs) rapidly mobilize innate and adaptive immunity to achieve antiviral status in the lung^42^. As shown in **Figure 4g**, RBD monomers did not induce apparent TRMs in the lung tissues, however the ICO-RBD nanovaccines generated 1.5-fold more TRMs in the lungs after boost immunization. Together, these results indicate that the ICO-RBD nanovaccines were able to induce marked T cell immune responses in addition to the B cell responses.

**Figure 4.**
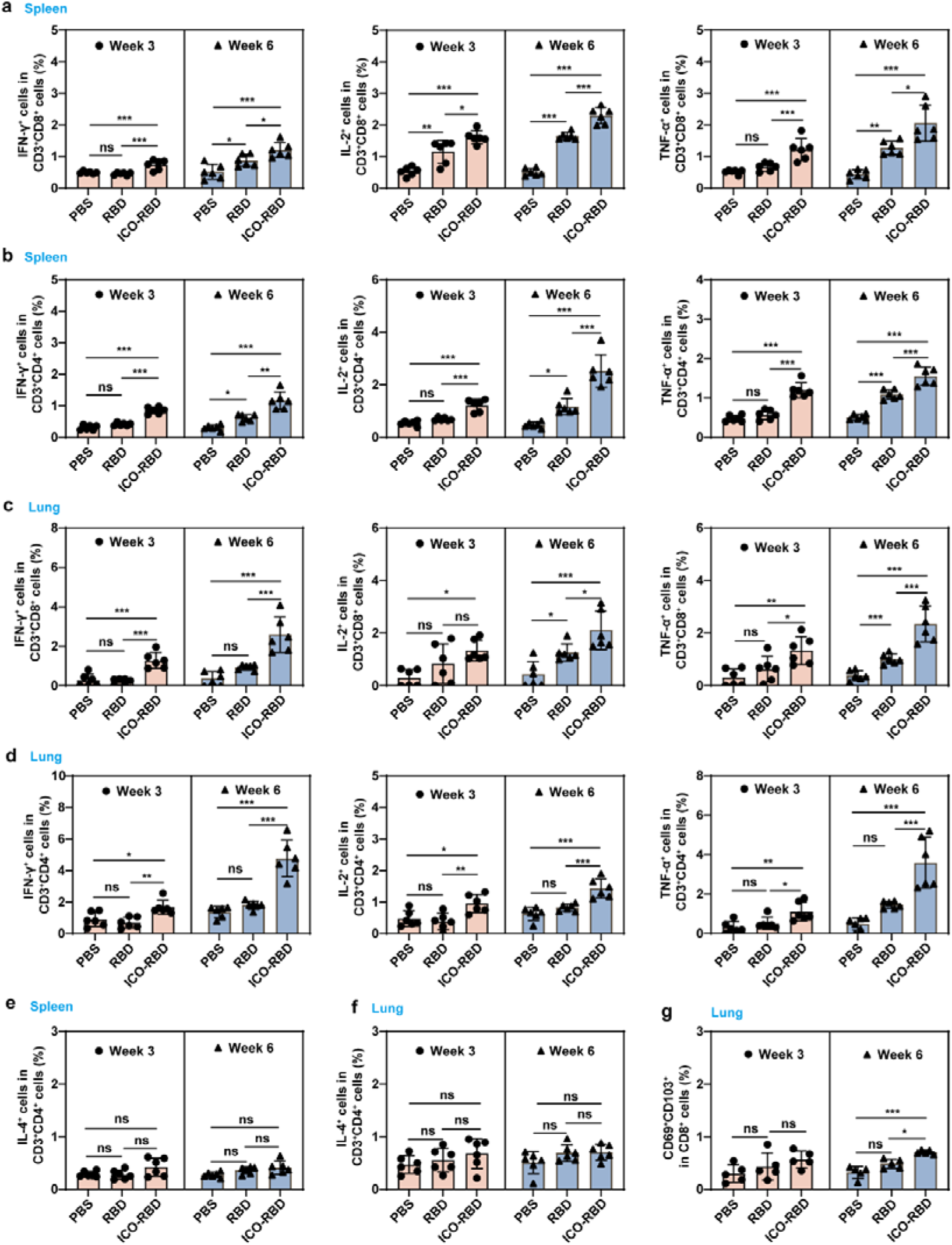
T cell responses induced by ICO-RBD nanovaccines in mice. The mice were randomly divided into 3 groups and priming- and boost-vaccinated with the indicated vaccines at week 0 and 3, respectively. The spleens and lungs of mice were harvested at week 3 (before boost immunization) and 6. The harvested cells were stimulated with SpyCatcher-RBD proteins overnight. **(a-b)** Proportions of IFN-γ^+^/IL-2^+^/TNF-α^+^ cells in CD3^+^CD8^+^ T cells (a) and CD3^+^CD4^+^ T cells (b) within the spleen, as detected using intracellular cytokine staining assays and flow cytometry (n = 6). **(c-d)** Proportions of IFN-γ^+^/IL-2^+^/TNF-α^+^ cells in CD3^+^CD8^+^ T cells (c) and CD3^+^CD4^+^ T cells (d) within the lung, as detected using intracellular cytokine staining assays and flow cytometry (n = 6). **(e-f)** Proportions of IL-4^+^ cells in CD3^+^CD4^+^ T cells within the spleen (e) and lung (f), as detected using intracellular cytokine staining assays and flow cytometry (n = 6). **(g)** Percentages of CD69^+^CD103^+^ TRMs in CD8^+^ T cells in the lung, as detected using flow cytometry (n = 6). The data were processed on GraphPad Prism 8 and are presented as the mean ± SD. Statistical significance (*P* value) was calculated by one-way ANOVA followed by Tukey’s test. *, *P* < 0.05; **, *P* < 0.01; ***, *P* < 0.001. ns, *P* > 0.05, no significant difference.

### ICO-RBD nanovaccine induction of antigen presentation and germinal center responses in drainage lymph nodes

To further elucidate how ICO-RBD nanovaccines are recognized and processed by the host immune system, and participate in immune responses, we injected (intramuscular) mice with Cy5.5-labeled RBD monomers or ICO-RBD nanovaccines (ICO-RBD-6c×5-10 nm) and isolated the inguinal lymph nodes after different time intervals (up to 14 days). As shown in **Figure 5a**, compared with RBD monomers, an increased amount of the ICO-RBD nanovaccine accumulated in the inguinal lymph nodes. A fraction of ICO-RBD nanovaccines was still present in the inguinal lymph nodes until day 7 (**Figure 5a**), which facilitated long-term immune stimulation of B and T cells.

**Figure 5.**
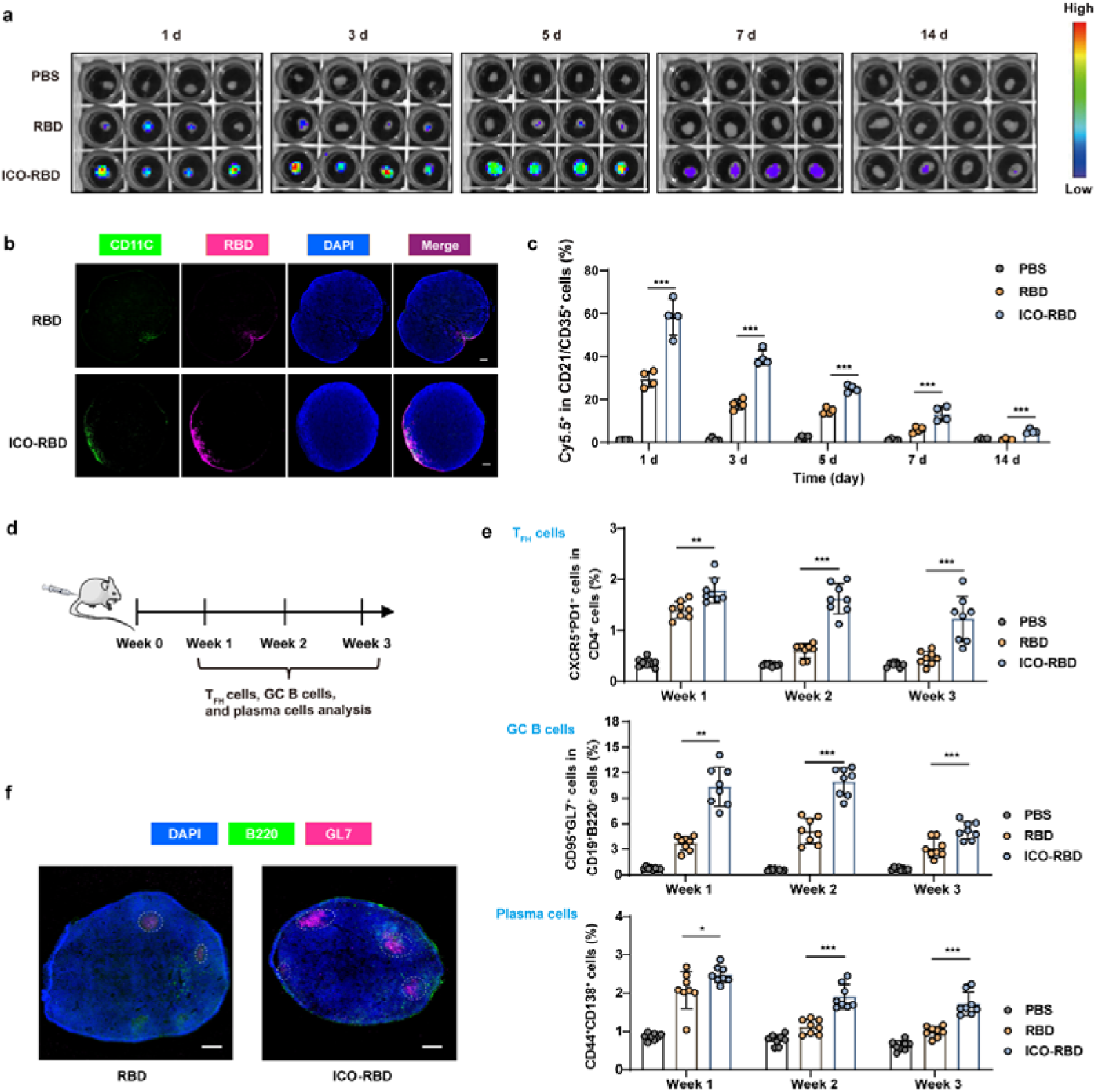
Antigen presentation and germinal center responses in drainage lymph nodes induced by ICO-RBD nanovaccines. **(a)** ICO-RBD nanovaccine accumulation in lymph nodes. BALB/c mice were intramuscularly immunized with Cy5.5-labelled RBD monomers (SpyCatcher-RBD proteins) or Cy5.5-labelled ICO-RBD nanovaccines. After different time intervals, the inguinal lymph nodes were obtained and then fluorescence imaging was performed. **(b)** Immunofluorescence detection of inguinal lymph nodes on day 1 and 3. Green, CD11C^+^ DCs, Red, Cy5.5-labelled RBD. Scale bar, 200 μm. **(c)** Percentages of CD21/CD35^+^Cy5.5^+^ follicular DCs in CD11C^+^ DCs in the inguinal lymph nodes after different time up to 14 days, as detected using flow cytometry (n = 4). **(d)** Schematic illustration of the experiment schedule to analyze germinal center responses. BALB/c mice were randomly divided into PBS, RBD and ICO-RBD groups. The inguinal lymph nodes were harvested at week 1, 2 and 3 after priming immunization. **(e)** Percentages of T_FH_ cells (CXCR5^+^PD1^+^) in CD4^+^ cells, GC B cells (CD95^+^GL7^+^) in CD19^+^B220^+^ cells and plasma cells (CD44^+^CD138^+^), as detected using flow cytometry (n = 8). **(f)** Immunofluorescence detection of inguinal lymph nodes on day 7. The white circles indicate the germinal centers, in which there are B220^+^GL7^+^ GC B cells. Scale bar, 200 μm. The data were processed on GraphPad Prism 8 and are presented as the mean ± SD. Statistical significance (*P* value) was calculated by one-way ANOVA followed by Tukey’s test. *, *P* < 0.05; **, *P* < 0.01; ***, *P* < 0.001.

Immunofluorescence detection of inguinal lymph nodes showed that the ICO-RBD nanovaccines and dendritic cells (DCs) were obviously co-localized (**Figure 5b**). A detailed analysis of vaccine uptake by antigen-presenting cells indicated that DCs (CD11C^+^), mature DCs (CD11C^+^CD80^+^ and CD11C^+^CD86^+^) and macrophages (F4/80^+^) captured 1.6, 1.8, 1.9 and 2.2 times, respectively, amount of ICO-RBD nanovaccines compared to RBD monomers (**Supplementary Figures 6a-d**). There were no significant differences in the amounts of ICO-RBD nanovaccine or RBD monomers captured by CD103^+^ migrated DCs (CD11C^+^CD103^+^) (**Supplementary Figure 6e**). However, differences were evident in the CD103^-^ resident DCs (CD11C^+^CD103^-^), as well as in the follicular DCs (CD11C^+^CD21/CD35^+^; **Supplementary Figures 6f-g**). Long-term monitoring of follicular DCs showed that ICO-RBD nanovaccines existed in follicular DCs for a longer time than RBD monomers (**Figure 5c**), which was conducive for follicular DCs to display the antigens on the surface and present them to B cells^43^. Thus, the ICO-RBD nanovaccines actively drained into the lymph nodes, and were more easily captured by resident and follicular DCs than monomeric RBD, which may contribute to the enhanced B and T cell immunity induced by the ICO-RBD nanovaccines.

Nanoparticle vaccines are more easily captured by DCs and macrophages than soluble vaccines, thus promoting the coordination of T follicular helper cells (T_FH_) and B cells, which is required for a prolonged and effective vaccination to drive antibody-mediated humoral immunity^44, 45^. To evaluate the germinal center responses, we measured T_FH_ cells (CD4^+^PD1^+^CXCR5^+^), germinal center (GC) B cells (CD19^+^B220^+^GL7^+^CD95^+^) and plasma cells (CD44^+^CD138^+^) in the inguinal lymph nodes at week 1, 2 and 3 after the priming immunization (**Figure 5d**). Although RBD monomers evoked measurable T_FH_ cell, GC B cell and plasma cell responses, induction with the ICO-RBD nanovaccines elicited respective responses in these cell populations that were 2.7, 2.1 and 1.7 times greater at week 2 (**Figure 5e**). Importantly, the ICO-RBD nanovaccines maintained elevated frequencies of T_FH_ cells, GC B cells and plasma cells for a longer period than RBD monomers, as indicated by the 2.7-, 1.7- and 1.7-fold greater responses of these cells at week 3 (**Figure 5e**). The serum titers of RBD-specific IgG in the ICO-RBD nanovaccine group gradually increased from week 1 to week 3, while the serum titers in the RBD group remained stable (**Supplementary Figures 7a-b**). On day 7, there were more germinal centers in lymph nodes in the ICO-RBD nanovaccine group than that in the RBD group (**Figure 5f**). Taken together, our data demonstrate that the ICO-RBD nanovaccines were more effectively captured by antigen-presenting cells and stimulated stronger germinal center responses than RBD monomers, thus enabling more effective antigen presentation, T-B crosstalk and B cell maturation.

### Comparison of ICO-RBD nanovaccines and trimeric mRNA vaccines containing Omicron RBD

To verify the compatibility of ICO scaffold for different antigens, we constructed ICO-RBD-6c×5-10 nm nanovaccines containing Omicron RBD (ICO-RBD_omic_). We used the “engraving-printing” strategy to decorate the RBD proteins of Omicron [B.1.1.529] onto ICO, except that the part responsible for the connection of RBD to polyT changed from SpyCatcher-SpyTag ligation to His-Nitrilotriacetic acid (NTA) affinity. For trimeric mRNA vaccine containing Omicron RBD (mRNA-RBD_omic_), we transcribed the RBD mRNA of Omicron [B.1.1.529] containing the C-terminal T4 fibritin foldon trimerization domain *in vitro*, and adopted the same lipid nanoparticle formulation as that of BNT162b2 to deliver mRNA and prepare mRNA-RBD ^46^.

Mice were immunized with two intramuscular injections of RBD_omic_, mRNA-RBD_omic_ and ICO-RBD_omic_, respectively, formulated with AddaVax adjuvant (**Figure 6a**). Both mRNA-RBD_omic_ vaccines and ICO-RBD_omic_ nanovaccines stimulated similar levels of IgG titers, both dramatically higher than those in the RBD_omic_ group, either at week 2 after priming immunization or at week 5 after boost immunization (**Figures 6b-c**). In terms of broad spectrum, the IgG in both mRNA-RBD_omic_ and ICO-RBD_omic_ groups were able to recognize RBD of all 11 variants we examined (**Figure 6d-e**). However, the ICO-RBD_omic_ nanovaccines activated more IL-2^+^CD4^+^ and TNF-α^+^CD8^+^ T cells than the mRNA-RBD_omic_ vaccines (**Figures 6f-h**). In addition, there was no apparent TRMs in the lung tissues in the mRNA-RBD_omic_ group, and the ICO-RBD_omic_ nanovaccines generated 3.3-fold more TRMs in the lungs after boost immunization (**Figure 6i**). In general, compared with the trimeric mRNA vaccines, the ICO-RBD nanovaccines stimulated similar humoral immune effects and exhibited a slight advantage in T cell immune activation.

**Figure 6.**
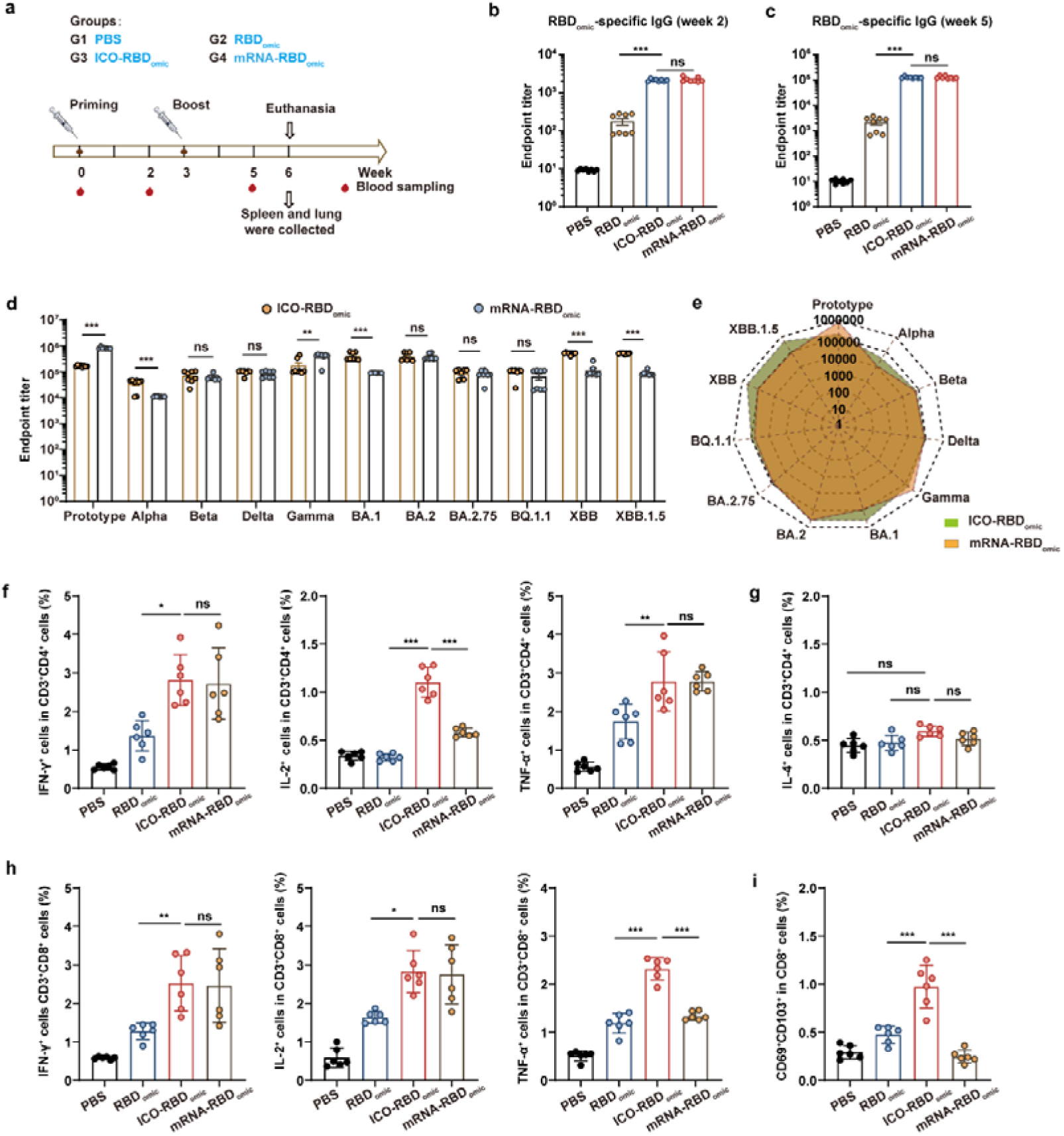
Comparison of ICO-RBD nanovaccines and trimeric mRNA vaccines containing Omicron RBD. **(a)** Schematic illustration of the vaccination schedule. Mice were randomly divided into 4 groups: PBS (control), Omicron RBD monomers (RBD_omic_), trimeric mRNA vaccine containing Omicron RBD (mRNA-RBD_omic_) and ICO-RBD-6c×5-10 nm nanovaccines containing Omicron RBD (ICO-RBD_omic_). Mice were intramuscularly immunized with two doses of 50 μL vaccines at week 0 and 3, mixed with 50 μL AddaVax. **(b)** Omicron RBD-specific IgG titers in serum, measured at week 2 (b) and 5 (c) by ELISA (n = 8). **(c-d)** Serum titers of IgG against variants, measured at week 5 by ELISA (n = 8). Different RBD proteins were used to coat 96-well ELISA plates, including prototype (Wuhan-Hu-1 strain), Alpha [B.1.1.7], Beta [B.1.351], Gamma [P.1], Delta [B.1.617.2] and Omicron [BA.1, BA.2, BA.2.75, BQ.1.1, XBB and XBB.1.5]. **(f)** Proportions of IFN-γ^+^/IL-2^+^/TNF-α^+^ cells in CD3^+^CD4^+^ T cells within the spleen, as detected at week 6 using intracellular cytokine staining assays and flow cytometry (n = 6). **(g)** Proportions of IL-4^+^ cells in CD3^+^CD4^+^ T cells within the spleen, as detected at week 6 using intracellular cytokine staining assays and flow cytometry (n = 6). **(h)** Proportions of IFN-γ+/IL-2+/TNF-α+ cells in CD3^+^CD8^+^ T cells (d) and CD3^+^CD4^+^ T cells within the spleen, as detected at week 6 using intracellular cytokine staining assays and flow cytometry (n = 6). **(i)** Percentages of CD69^+^CD103^+^ TRMs in CD8^+^ T cells in the lung, as detected at week 6 using flow cytometry (n = 6). The data were processed on GraphPad Prism 8. The data are presented as the mean ± SEM in panel b, c and d, and the mean ± SD in panels f-i. Statistical significance (*P* value) was calculated by one-way ANOVA followed by Tukey’s test. *, *P* < 0.05; **, *P* < 0.01; ***, *P* < 0.001. ns, *P* > 0.05, no significant difference.

Compared to subunit protein vaccines, the non-viral vector-delivered nucleic acid-based vaccines mimics infection or immunization with live microorganisms. Therefore, the nucleic acid-based vaccines can generate cell-mediated immunity at the same time as humoral immunity, and the subunit protein vaccines require adjuvants to activate effective cell-mediated immunity. However, our ICO-displayed subunit nanovaccines activated robust cell-mediated immunity, even slightly higher than the mRNA vaccines. Considering that ICO was derived from the assembly of genomic DNA of M13mp18 bacteriophage, ICO not only has the function of a scaffold to display subunit antigens, but also may have the function of an adjuvant similar to nucleic acid-based vaccines to mimics viral invasion, which is a potential advantage of the DNA-based nanocarriers.

### Immunogenicity of ICO scaffold and biosafety of ICO-RBD nanovaccines

To analyze the immunogenicity of ICO *in vitro* and *in vivo*, bone marrow-derived dendritic cells (BMDCs) were isolated and treated with different concentrations of ICO. As shown in **Figure 7a**, ICO can stimulate BMDCs’ maturation in a dose-dependent manner, indicated by the gradual increase in CD80^+^CD86^+^ BMDCs. Even after decoration of RBD, ICO-RBD still possessed the ability to stimulate BMDCs’ maturation (**Figure 7a**). After intramuscular injection, ICO also generated an obvious elevation of CD80^+^CD86^+^ DCs in the drainage lymph node (**Figure 7b**). These results suggest that ICO are inherently immunogenic.

**Figure 7.**
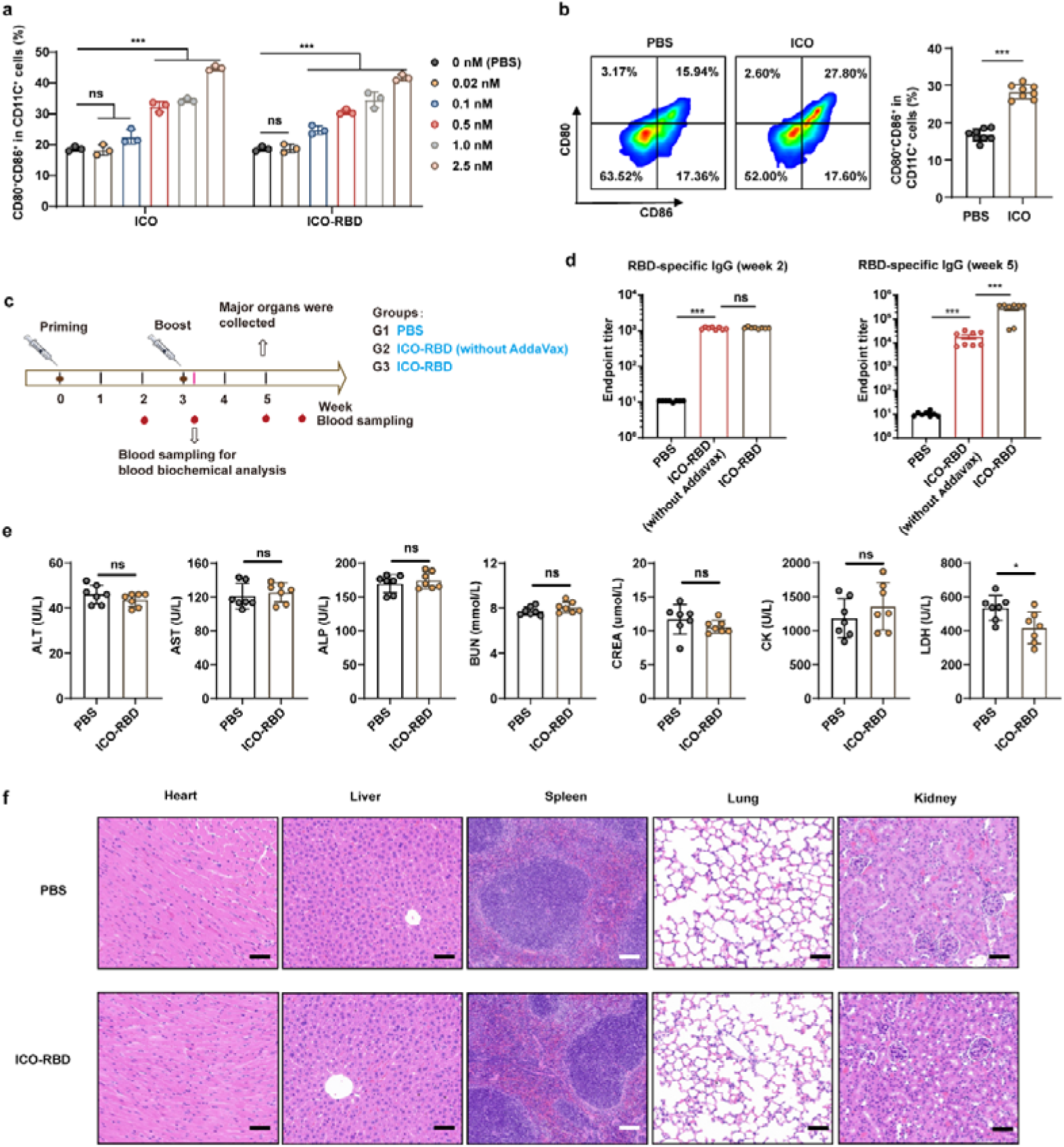
Immunogenicity of ICO scaffold and biosafety of ICO-RBD nanovaccines. **(a)** Percentages of CD80^+^CD86^+^ in CD11C^+^ cells in BMDCs after treatment with different concentrations of ICO for 12 h, as detected using flow cytometry (n = 3). **(b)** Percentages of CD80^+^CD86^+^ in CD11C^+^ DCs in the inguinal lymph nodes, as detected using flow cytometry (n = 8). Mice were intramuscularly injected with two doses of ICO at week 0 and 3, and the inguinal lymph nodes were harvested after 3 weeks. **(c)** Schematic illustration of the vaccination schedule. Mice were randomly divided into 3 groups: PBS (control), ICO-RBD nanovaccines without AddaVax (ICO-RBD without AddaVax), and ICO-RBD nanovaccines with AddaVax (ICO-RBD vaccines). Mice were intramuscularly immunized with two doses of ICO-RBD nanovaccines at week 0 and 3, mixed with or without 50 μL AddaVax. **(d)** RBD-specific IgG titers in serum, measured at week 2 and 5 by ELISA (n = 8). **(e-f)** Blood biochemical analysis (e, n = 7) at day 22 and histological evaluation of major organs (f) at week 5. Black scale bar, 50 μm, white scale bar, 100 μm. Mice were intramuscularly immunized with two doses of ICO-RBD nanovaccines at week 0 and 3, mixed with 50 μL AddaVax. The data were processed on GraphPad Prism 8. The data are presented as the mean ± SEM in panel d, and the mean ± SD in panel a, b and e. Statistical significance (*P* value) was calculated by one-way ANOVA followed by Tukey’s test. *, *P* < 0.05; ***, *P* < 0.001. ns, *P* > 0.05, no significant difference.

Subsequently, we assessed whether the immunogenicity of ICO could replace adjuvants in the vaccination. Mice received two intramuscular injections of ICO-RBD nanovaccines and ICO-RBD nanovaccines without AddaVax adjuvant (ICO-RBD without AddaVax), respectively (**Figure 7c**). Although the IgG titers in the ICO-RBD and ICO-RBD without AddaVax groups were similar after priming immunization, the IgG titers in the ICO-RBD group were significantly higher than those in the ICO-RBD without AddaVax group after boost immunization (**Figure 7d**). Therefore, the additional adjuvant is still necessary for our ICO-based nanovaccines.

Finally, we evaluated the biosafety of ICO-RBD nanovaccines. Two intramuscular injections of ICO-RBD nanovaccines did not induce significant changes in blood biochemical analysis (**Figure 7e**), nor did they cause obvious histological abnormalities in major organs (**Figure 7f**).

## Conclusion

In summary, we employed an icosahedral DNA origami, ICO, to construct tailor-made nanovaccines with shapes and sizes closely resembling SARS-CoV-2, as an RBD antigen display platform. Based on an “engraving-printing” strategy that can efficiently and specifically link site-modified RBD antigens onto the surface of ICO, we achieved a rational design and accurate assembly of SARS-CoV-2 RBD nanovaccines. We also scrutinized several parameters of the surface antigen pattern, including antigen spacing, antigen copies within clusters and cluster numbers, for their influence on B cell activation. Finally, we systematically evaluated the immunogenicity of the optimized nanovaccines, ICO-RBD-6c×5-10 nm, which evoked more robust and enduring humoral and T cell immune responses than soluble RBD antigens in mice. The ICO scaffold and the optimized surface antigen pattern were also suitable for the display of Omicron RBD, and the constructed nanovaccines exhibited the comparable immune effects as the mRNA vaccines. The structure-function relationship between antigen configuration and immunogenicity that we demonstrate through the use of the DNA origami platform may pave the way for the future development of potent vaccines for SARS-CoV-2 and/or other viruses.

## Materials and methods

### Reagents and materials

Oligonucleotides, purified by polyacrylamide gel electrophoresis (PAGE), used for folding ICO were synthesized by Beijing Ruibio Biotech Co., Ltd (Beijing, China). The genomic DNA of M13mp18 bacteriophage was obtained from Beijing Intell Nanomedicine (Beijing, China). DBCO-polyT (20 × T), SpyTag-N3, Cy5.5-SpyTag and FITC-SpyTag were synthesized by China Peptides Co., Ltd. (Shanghai, China). The RBD-specific BCR expression plasmid, pRRL EuB29 COVA2-15 IgGTM. BCR. GFP. WPRE was a kind gift from Drs. Marit J. van Gils and Rogier W. Sanders of the Department of Medical Microbiology at the University of Amsterdam. The SARS-CoV-2 pseudovirus neutralization test kit (Cat: #L02087A), including the SARS-CoV-2 pseudovirus Luc Reporter and HEK-293 cells expressing hACE2, was purchased from GenScript (Nanjing, China).

The antibodies used for flow cytometry were as follows: anti-His Tag APC (BioLegend, USA, Cat: #362605), anti-human IgG (G18-145) APC (BD, USA, Cat: #550931), anti-mouse CD3 APC (BioLegend, Cat: #100312), anti-mouse CD19 APC (BioLegend, Cat: #115512), anti-mouse CD138 APC (Biolegend, Cat: #142506), anti-mouse CXCR5 APC (Biolegend, Cat: #145506), anti-mouse CD103 APC (Biolegend, Cat: #121414), anti-mouse CD8 FITC (Biolegend, Cat: #100706), anti-mouse CD11C FITC (Biolegend, Cat: #117306), anti-mouse CD95 (Fas) FITC (Biolegend, Cat: #152606), anti-mouse IgG FITC (Biolegend, Cat: #406001), anti-mouse CD4 PE (Invitrogen, USA, Cat: #12-0042-85), anti-mouse CD44 PE (Biolegend, Cat: #103008), anti-mouse CD80 PE (Biolegend, Cat: #104708), anti-mouse CXCR3 PE (Biolegend, Cat: #126505), anti-mouse CD103 PE (Biolegend, Cat: #121406), anti-mouse CD21/CD35 PE (Biolegend, Cat: #123410), anti-mouse CD69 PE (Biolegend, Cat: #104508), anti-mouse CD44 PE (Biolegend, Cat: #103008), anti-mouse CD86 PE/Cyanine7 (Biolegend, Cat: #105014), anti-mouse IFNγ PE/Cyanine7 (Biolegend, Cat: #505826), anti-mouse F4/80 PE/Cyanine7 (Biolegend, Cat: #123114), anti-mouse CD279 (PD-1) PE/Cyanine7 (Biolegend, Cat: #109109), anti-mouse/human GL7 PE/Cyanine7 (Biolegend, Cat: #144619), anti-mouse EpCAM PE/Cyanine7 (Biolegend, Cat: #118215), anti-mouse CD38 PE/Cyanine7 (Biolegend, Cat: #102718), anti-mouse IL2 PE/Cyanine7 (Biolegend, Cat: #503831), anti-mouse IL4 PE/Cyanine7 (Biolegend, Cat: #504117), anti-mouse TNF-α PE/Cyanine7 (Biolegend, Cat: #506323), anti-mouse CD62L PE/Cyanine7 (Biolegend, Cat: #104418), anti-mouse CD8a PE/Cyanine7 (Biolegend, Cat: #100722), anti-mouse/human B220 APC/Cyanine7 (Biolegend, Cat: #103223), anti-mouse CD4 PerCP (Biolegend, Cat: #100432).

The antibodies used for immunofluorescent staining were as follows: Alexa Fluor™ 488 conjugated CD11C (N418) antibody (eBioscience, USA, Cat: 53-0114-82), anti-mouse Alexa Fluor™ 488 conjugated B220 (RA3-6B2) antibody (eBioscience, Cat: 53-0452-82), Alexa Fluor® 647 anti-mouse/human GL7 antigen (Biolegend, Cat:144605)

### Protein expression and purification

The expression and purification of SpyCatcher-RBD (residues 319-541, prototype) from HEK-293 cells were completed by AtaGenix Laboratories Co., Ltd. (Wuhan, China.). Briefly, the 8 × His-labelled SpyCatcher-RBD DNA sequence (**Supplementary Figure 2a**) was cloned into the pATX2 vector and transfected into HEK-293 cells. After 7 days, the culture medium soluble fraction was collected, and the His-labelled target proteins were purified by Ni-NTA affinity, and then concentrated after elution with Tris-buffer containing imidazole. The purified His-labelled proteins were identified by Coomassie blue staining and western blot analysis. The protein concentration was determined by the BCA method. His-labelled Omicron RBD (residues 319-541, B.1.1.529) (His-Omicron RBD) and His-labelled RBD proteins of different variants were obtained using a similar method.

### ICO folding and purification

ICO design (115.5 bp edge length) was performed using the algorithmic framework software, DAEDALUS (http://daedalus-dna-origami.org). The sequence information of the staple strands for ICO fabrication are provided in **Supplementary Data 1**. The sequence information of the capture strands in the different ICO-RBD nanovaccines is also shown in **Supplementary Data 1**. DNA origami was fabricated by the following one-pot reaction: 20 nM M13mp18 scaffold was mixed with a 100 nM staple or capture strand mixture in 1 × TAE-Mg^2+^ buffer (40 mM Tris, 20 mM acetic acid, 2 mM EDTA, 12.5 mM MgCl_2_, pH 8.0). The mixture was then self-assembled in a thermal cycler (T100, Bio-Rad) through the following program: 95°C for 5 min, 80-75°C at 1°C per 5 min, 75-30°C at 1°C per 20 min and 30-25°C at 1°C per 10 min. The freshly prepared DNA origami was purified using 100 KDa MWCO centrifugal filters (Merck Millipore, UFC5100BK, USA) for 3 rounds to remove excess staple and capture strands. The concentration of the origami preparation was quantified using a NanoDrop 2000 (Thermo Fisher Scientific, USA) and stored at 4□ for subsequent analysis. The origami was characterized by electrophoresis in a 0.6% agarose gel, and was imaged by TEM (JEM-1400) or AFM (Bruker, Multimode 8).

### Antigen attachment to ICO

For SpyCatcher-RBD (prototype) protein attachment, SpyTag-N3 and DBCO-polyT were mixed at a molar ratio of 1:1, and reacted overnight at 4□ to obtain SpyTag-polyT conjugates. Purified ICO was mixed with SpyTag-polyT conjugates at a molar ratio of 5:1 of poly T: polyA on the ICO.

PolyT-polyA hybridization was performed by the following thermal cycler program: 45-25□ at 1°C per 5 min for 5 cycles, and a hold at 4□. The products (ICO-SpyTag) were purified using 100 KDa MWCO centrifugal filters (Merck Millipore, UFC5100BK,) for 3 rounds to remove excess SpyTag-polyT conjugates. SpyCatcher-RBD proteins were then added to ICO-SpyTag at a molar ratio of 3:1 of SpyCatcher: SpyTag. This mixture was incubated at room temperature for 3 h, and excess Spycatcher-RBD proteins were removed by ultrafiltration.

For His-Omicron RBD attachment, NH_2_-PolyT (100 μM) and NTA-PEG_1000_-NHS (3 mM) were shaken at 25 °C, 400 rpm for 2 h. The pre-cooled ethanol (90%) and potassium acetate (pH5.5, 60 μM) were mixed, and left at -80□ for 30 min, then 14000 g and centrifuged at 4□ for 30 min. The supernatant was removed. After re-suspension with 100 μl PBS (pH7.4), the above ethanol precipitation steps were repeated. The supernatant was discarded and dried, and the obtained NTA-polyT was dissolved by TE Buffer for subsequent use. Purified ICO was mixed with NTA-polyT conjugates at a molar ratio of 5:1 of polyT: polyA on the ICO. PolyT-polyA hybridization was performed by the following thermal cycler program: 45-25□ at 1°C per 5 min for 5 cycles, and a hold at 4□. The products (ICO-NTA) were purified using 100 KDa MWCO centrifugal filters (Merck Millipore, UFC5100BK, USA) for 3 rounds to remove excess NTA-polyT conjugates. His-Omicron RBD proteins were then added to ICO-NTA at a molar ratio of 3:1 of His: NTA. This mixture was incubated at room temperature for 2 h, and excess His-Omicron RBD proteins were removed by ultrafiltration.

### Protein quantification of ICO coverage with antigen

The antigen loading efficiency was quantified by ratio absorbance measurement. Specifically, the concentration of the DNA origami was measured by absorbance using a NanoDrop 2000 spectrophotometer at 260 nm, and the extinction coefficient of 10.9 × 10^7^ M^−1^ cm^−1^ was used for the calculation. The concentration of antigens was quantitatively determined using the BCA standard curve method. The antigen loading efficiency was defined as the percentage of measured protein concentration in nanovaccines compared to the theoretical concentration in the nanovaccines. At least 3 independent experiments were performed.

### Preparation of mRNA vaccines

mRNA was synthesized through *in vitro* transcription using the HiScribe T7 mRNA Kit with CleanCap Reagent AG (NEB, China, Cat: E2080S). The templates for *in vitro* transcription were linearized plasmids, containing a T7 promoter, 5’-end UTR, signal peptide, Omicron RBD (residues 319-541, B.1.1.529), linker, T4 fibritin foldon trimerization domain, stop codon, 3’-end UTR and 120 polyA tail. The production of lipid nanoparticles was carried out using the NanoAssemblr Benchtop microfluid system (Precision NanoSystems, Canada). The lipid nanoparticle formulation was same as that of BNT162b2: ALC-0315: DSPC: Cholesterol: ALC-0159 = 46.3: 9.4: 42.7: 1.6.

### Generation of B-RBD cells

Ramos B cells were obtained from Cell Resource Center, Institute of Basic Medicine, Chinese Academy of Medical Sciences, and were authenticated by the short tandem repeat analysis method and tested negative for mycoplasma contamination. The expression plasmid pRRL EuB29 COVA2-15 IgGTM. BCR. GFP. WPRE and the packaging plasmid mix (pMDL: pVSV-g: pRSV-Rev = 5: 3: 2) were co-transfected into HEK-293T cells at a ratio of 1:1 using lipofectamine 3000 (Invitrogen, USA, Cat: #L3000015). After 48 h, the supernatant from the cells was collected and filtered (0.45 μm), and then 50 µl of the concentrated viral supernatant was added to pre-plated Ramos B cells in 6-well plates (2 × 10^6^ cells/well). After 7 days of transfection, the GFP^+^IgG^+^ cell population (B-RBD cells) were sorted by FACS (BD Aria III).

### B cell binding assay

B-RBD cells were grown in RPMI 1640 medium (Gibco, Cat: #C11875500CP) supplemented with 10% fetal bovine serum (Gibco, Cat: #10091148) and 100 U/mL penicillin G sodium and 100 µg/mL streptomycin (Biological Industries, Cat: #03-031-1B). Cells (1 × 10^6^) were harvested, suspended in PBS and incubated with 0.5 μg/mL SpyCatcher-RBD proteins or different ICO-RBD nanovaccine formulations at the same protein concentration for 1, 2, 4, 6, 8 or 10 min at 4□. Next, the cells were washed three times in PBS and incubated with anti-His antibodies at 4°C for 40 min. The cells were then washed twice with PBS and SpyCatcher-RBD proteins or ICO-RBD nanovaccines bound to B-RBD cells were detected by flow cytometry. The gating strategies for flow cytometry analysis are shown in **Supplementary Figure 8**.

### B cell activation assay

B-RBD cells (1 × 10^7^ cells/mL) were combined with 4 μM Fluo-4 AM (Invitrogen, USA, Cat: #F14217) for 20 min at 37°C. The cells were then washed twice with HBSS (Solarbio, China, Cat: #H1025) and then incubated at 37°C for 30 min. The washed cells were resuspended in PBS, and the nanovaccine-induced intracellular calcium changes were detected by flow cytometry (excitation/emission: 495 nm/518 nm). After a 35 s fluorescence baseline recording, 50 μL SpyCatcher-RBD proteins or ICO-RBD nanovaccines (5 μg/mL RBD) were added to the cells at 2 × 10^6^ cells/mL and the fluorescence values were monitored for 3–5 min. The fluorescence value of a 50 μL PBS sample was used as a normalization standard at every time point.

### Animal vaccination

All animal experiments were performed in accordance with animal use protocols approved by the Committee for the Ethics of Animal Experiments, the Institutional Animal Care and Use Committee of the National Center for Nanoscience and Technology. Mice were housed at 20–22°C with a 12 h light/dark cycle and at 30–70% humidity. Female BALB/c mice (6 weeks old) were obtained from Charles River Laboratory Animal Technology Co. Ltd (Beijing, China).

For prototype RBD vaccine evaluation, six-week-old female BALB/c mice were randomly divided into 8 groups: PBS (control), ICO, RBD (SpyCatcher-RBD proteins), ICO + RBD (physical mixture), ICO-RBD-1c×5-10 nm, ICO-RBD-1c×5-40 nm, ICO-RBD-6c×1-10 nm and ICO-RBD-6c×5-10 nm. BALB/c mice were intramuscularly immunized with two doses of 50 μL nanovaccine containing 1 μg SpyCatcher-RBD at week 0 and 3, mixed with 50 μL AddaVax (InvivoGen, USA, Cat: #vac-adx-10). Serum samples were collected at week 2, 5, 8, 11 and 14, and RBD-specific IgG titers were measured by ELISA. Serum collected at week 5 and 14 was analyzed using a pseudovirus neutralization test. Spleens and lungs of mice were harvested at week 3 and 6 for T cell analysis. All mice were euthanized at week 14 and memory immune cells were evaluated.

For comparison of ICO-RBD nanovaccines and trimeric mRNA vaccines containing Omicron RBD, mice were randomly divided into 4 groups: PBS (control), RBD_omic_, mRNA-RBD_omic_ and ICO-RBD_omic_. The RBD_omic_ and ICO-RBD_omic_ vaccines contained 2 μg His-Omicron RBD proteins, and the mRNA-RBD_omic_ vaccines contained 10 μg mRNA. Mice were intramuscularly immunized with two doses of 50 μL vaccines at week 0 and 3, mixed with 50 μL AddaVax. Serum samples were collected at week 2 and 5, and RBD-specific IgG titers were measured by ELISA. Spleens and lungs of mice were harvested at week 6 for T cell analysis. All mice were euthanized at week 14 and memory immune cells were evaluated.

### ELISA

SpyCatcher-RBD (prototype), His-Omicron RBD proteins or different RBD proteins of variants were diluted to 5 μg/mL with ELISA coating buffer (Solarbio, Cat: #C1055) and used to coat 96-well ELISA plates (Thermo Fisher, USA, Cat: #442404) at 4□ overnight. The washed plates were blocked with 5% skim milk in PBS for 2 h. The serum was serially diluted and added to each well, and incubated at room temperature for 2 h. After washing the plates 3 times with PBS containing 1% Tween-20 (PBST), a goat anti-mouse IgG HRP antibody (Proteintech, USA, Cat: #SA00001-1) was diluted 1: 5000 in 5% skim milk, and added to the plates, which were then incubated at 37□ for 1 h. After washing with PBST, the plates were developed with 3,3’,5,5’-tetramethylbenzidine (TMB; Solarbio, Cat: #PR1200). The color reaction was stopped using ELISA stop solution (Solarbio, Cat: #C1058), and the absorbance at 450 nm was read in a microplate reader (BioTEK, USA, Synergy H1). The data were analyzed by nonlinear regression to calculate the endpoint titer.

### Pseudovirus neutralization assay

The pseudovirus neutralization assay was performed according to the manufacturer’s instructions. Briefly, the serum samples and positive control (ACE2-Fc proteins) were diluted in a continuous gradient, and then mixed with resuscitated SARS-CoV-2 pseudovirus in a volume of 1: 1 at room temperature for 1 h, where the virus titer was 3 × 10^4^ TCID_50_/mL. The mixture was added to HEK-293 cells expressing hACE2 (6 × 10^5^/mL) and incubated in a 5% CO_2_ incubator at 37□ for 48 h. Luciferase activity was analyzed with a Luciferase Assay System (GenScript, Cat: #L00877C). The relative luminous unit (RLU) was standardized to the relative luminous unit of SARS-CoV-2 pseudovirus-infected cells in the absence of serum. Neutralization titer (ID_50_ value) was calculated as the serum dilution with 50% inhibition of infectivity.

### Memory T cells, antigen-specific MBCs and LLPC analysis

Splenocytes of mice were obtained and prepared as a single-cell suspension at week 14. For RBD-specific MBC analysis, Cy5.5-SpyTag was conjugated to SpyCatcher-RBD proteins to prepare “Cy5.5-RBD” fluorescent probes. Cells were incubated with Cy5.5-RBD for 1 h at 4□, washed 3 times with PBS, and then fixed and permeabilized. B220, IgG and CD38 antibodies were used to analyze antigen-specific MBCs in the splenocytes. The gating strategy for the flow cytometry analysis is shown in **Supplementary** Figure 9. For LLPC analysis, splenocytes and bone marrow cells (from the femur and tibia) were obtained, and red blood cells were removed, to obtain single-cell suspensions. FITC-RBD probes were prepared according to the above method. The cells were mixed with FITC-RBD for 1 h at 4□, and then washed 3 times with PBS. B220, CD138, EpCAM and CXCR3 antibodies were used to identify antigen-specific LLPCs. The gating strategy for the flow cytometry analysis is shown in **Supplementary** Figure 10. The central memory T cells and effector memory T cells in splenocytes were analyzed by flow cytometry after staining with CD3, CD8, CD44 and CD62L antibodies. The gating strategy for the flow cytometry analysis is shown in **Supplementary Figure 11**.

### Intracellular cytokine staining assays and TRM analysis

Spleens and lungs of mice were harvested at week 3 and 6. The spleens were ground and passed through a 40 μm cell strainer to obtain single cell suspensions. The lungs were cut into pieces and treated with freshly prepared digestion buffer (0.05% trypsin solution, 1 mg/mL DNase and 1 mg/mL collagenase D) for 30 min with shaking at 120 rpm in 37□. The cells were stimulated with SpyCatcher-RBD or His-Omicron RBD proteins overnight, and then the cells were collected and stained with CD3, CD4 and CD8 antibodies. Fixation and permeabilization were performed after surface marker staining, and the cells were sequentially stained with IFN-γ, IL-2, TNF-α and IL-4 antibodies. The proportion of IFN-γ^+^, IL-2^+^, TNF-α^+^ and IL-4^+^ cells in CD3^+^CD8^+^ T cells and CD3^+^CD4^+^ T cells were analyzed using flow cytometry. The gating strategies for the flow cytometry analysis are shown in **Supplementary Figure 12 and 13**. TRMs in the lung were analyzed after staining with CD8, CD69 and CD103 antibodies. The gating strategies for the flow cytometry analysis are shown in **Supplementary Figure 14**.

### DC and Macrophage studies

Six-week-old female BALB/c mice were intramuscularly immunized with 50 μL Cy5.5-labelled SpyCatcher-RBD proteins or Cy5.5-labelled nanovaccines containing 1 μg SpyCatcher-RBD proteins, mixed with 50 μL AddaVax. After different time, the inguinal lymph nodes were obtained and then fluorescence imaging was performed. Single cell suspensions were obtained after grinding the samples. The Cy5.5^+^ cells in DCs (CD11C^+^), resident DCs (CD103^-^CD11C^+^), migrated DCs (CD103^+^CD11C^+^), follicular DCs (CD21/CD35^+^CD11C^+^), mature DCs (CD11C^+^CD80^+^, CD11C^+^CD86^+^) and macrophages (F4/80^+^) were analyzed using flow cytometry. The gating strategies for the flow cytometry analysis are shown in **Supplementary Figure 15**. The inguinal lymph nodes were also collected on day 1 and 3, and the cryosections were cut to perform immunofluorescent staining.

### Analysis of GC B, T_FH_ and plasma cells

For GC B cells and T_FH_ cell response analysis, six-week-old female BALB/c mice were randomly divided into 3 groups: PBS, RBD (SpyCatcher-RBD proteins) and ICO-RBD nanovaccine. After priming immunization, the inguinal lymph nodes were harvested at week 1, 2 and 3. Single cell suspensions were then obtained by grinding and sieving the nodes. Next, the percentages of GC B cells (CD19^+^B220^+^CD95^+^GL7^+^), T_FH_ cells (CD4^+^CXCR5^+^PD-1^+^) and plasma cells (CD44^+^CD138^+^) were analyzed by flow cytometry. The gating strategies for the flow cytometry analysis are shown in **Supplementary Figure 16**. The inguinal lymph nodes were also collected on week 1, and the cryosections were cut to perform immunofluorescent staining.

### Culture of BMDCs

Briefly, bone marrow cells were flushed from the femurs and tibias of mice and cultured in RPMI 1640 supplemented with 10% FBS, 100 U/mL penicillin G sodium, 100 µg/mL streptomycin, 1% HEPES, 0.05 mM β-mercaptoethanol (β-ME), 20 ng/mL IL-4 (Cell Signaling Technology, USA, Cat: 5208SC) and 20 ng/mL GM-CSF (PeproTech, USA, Cat: 315-03-20UG), after lysed the red blood cells using ACK Lysis Buffer (Solarbio, China, Cat: R1010) to induce differentiation into immature BMDCs.

## Statistical analysis

Details on the statistical analysis for the experiments, including the number of samples, mean ± standard errors of the mean (SEM) or mean ± standard deviation (SD) are shown in the figure legends. Statistical analysis was performed using Prism 8 (GraphPad) on data pooled from at least 3 independent experiments. For multiple-group comparisons, one-way ANOVA followed by Tukey’s test, was applied.

## Acknowledgements

This work was supported by grants from the National Key R&D Program of China (2022YFB3808100 and 2021YFA0909900, X. Z.), Strategic Priority Research Program of Chinese Academy of Sciences (XDB36000000, G.N.), the CAS Project for Young Scientists in Basic Research (YSBR-010, X. Z.), the Beijing Natural Science Foundation (Z200020, X. Z.) and the National Natural Science Foundation of China (32222045 and 32171384, X. Z.).

## Conflicts of interest

Q. F., J. Z., Q. S. and X. Z. are inventors on a filed provisional application of Chinese patent (Nano-assembly of antigens based on DNA origami and the application in vaccine) submitted by the National Center for Nanoscience and Technology and Beijing Intell Nanomedicine that covers the potential diagnostic and therapeutic uses of DNA-antigen complex. The authors declare that they have no other competing interests.

## Author Contributions

Q. F., K. C., and L. Z. contributed equally to this work. Q. F. and X. Z. designed the research. Q. F., K. C., L. Z., X. G., J. L., G. L., N. M., C. X., M. T., L. C., X. W., J. Z., and Q. S. performed the research. All authors analyzed and interpreted the data. Q. F., G. N. and X. Z. wrote the paper. G. N. and X. Z. conceived and supervised the project.

## Data availability

The main data supporting the results in this study are available within the paper and its Supplementary Information/Data. There are no data from third-party or publicly available datasets. Other source data that support the findings of this study are available from the corresponding authors upon reasonable request.

