## Supplementary Figures 1-16 for "Rationally designed multimeric nanovaccines using icosahedral DNA origami for molecularly controlled display of SARS-CoV-2 receptor binding domain"

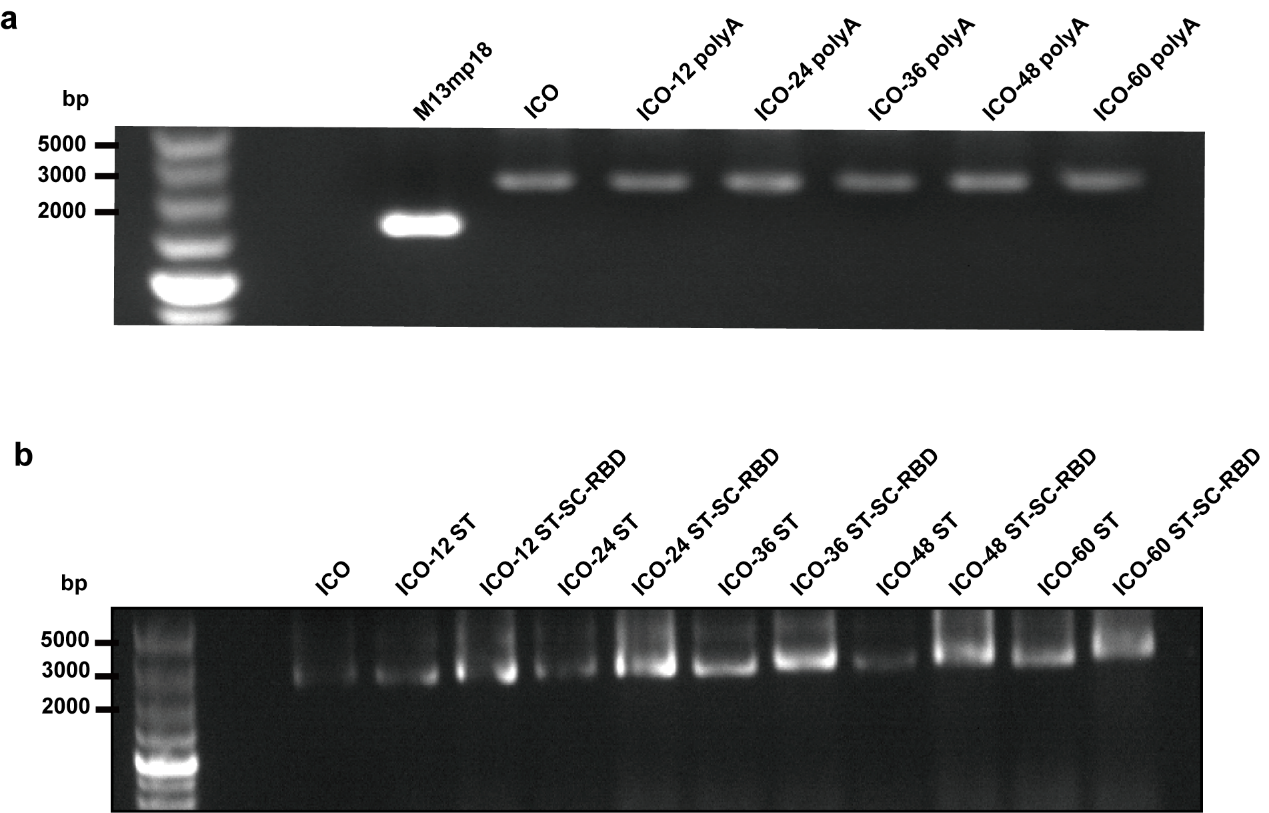

**Supplementary Figure 1.** Agarose gel electrophoresis of ICO. **(a)** Electrophoresis of ICO containing 12–60 capture strands with overhanging polyA. **(b)** Electrophoresis of ICO containing 12–60 SpyTag (ST) or SpyTag-SpyCatcher-RBD (ST-SC-RBD).

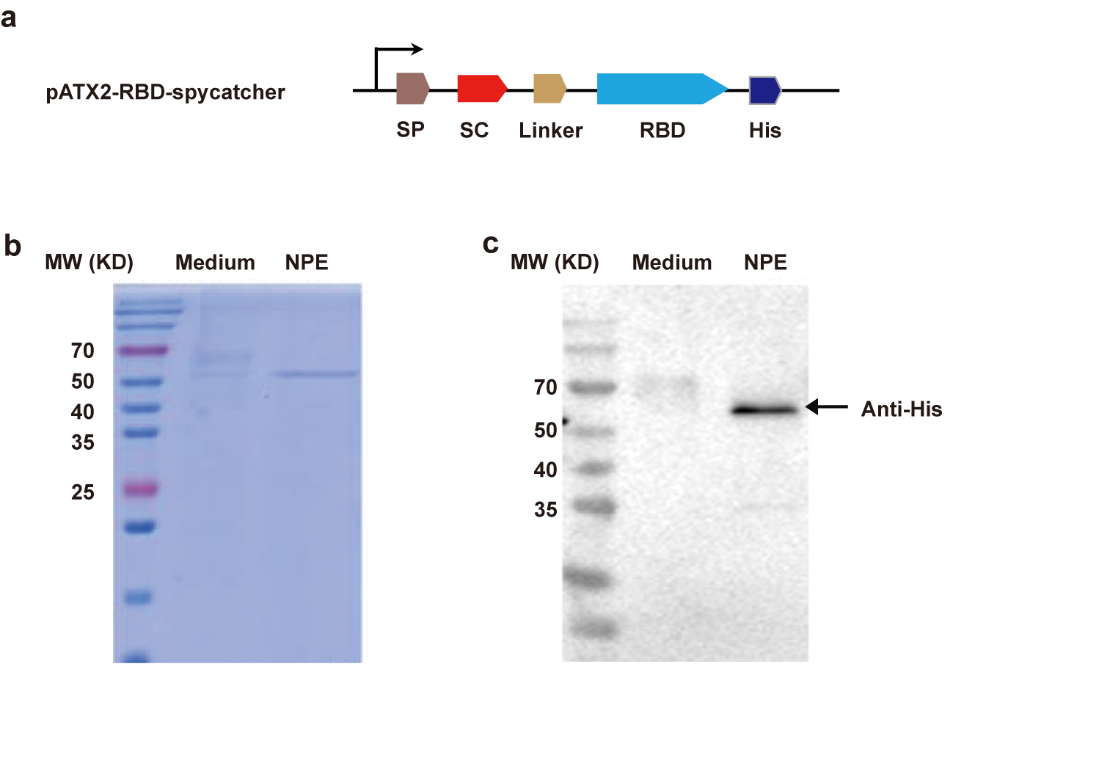

**Supplementary Figure 2. (a)** Schematic representation of the pATX2-SpyCatcher-RBD construct. **(b)** Coomassie blue staining and **(c)** western blot analysis of the fusion protein. SP, signaling peptide. SC, SpyCatcher. NPE, soluble fraction. MW, molecular weight.

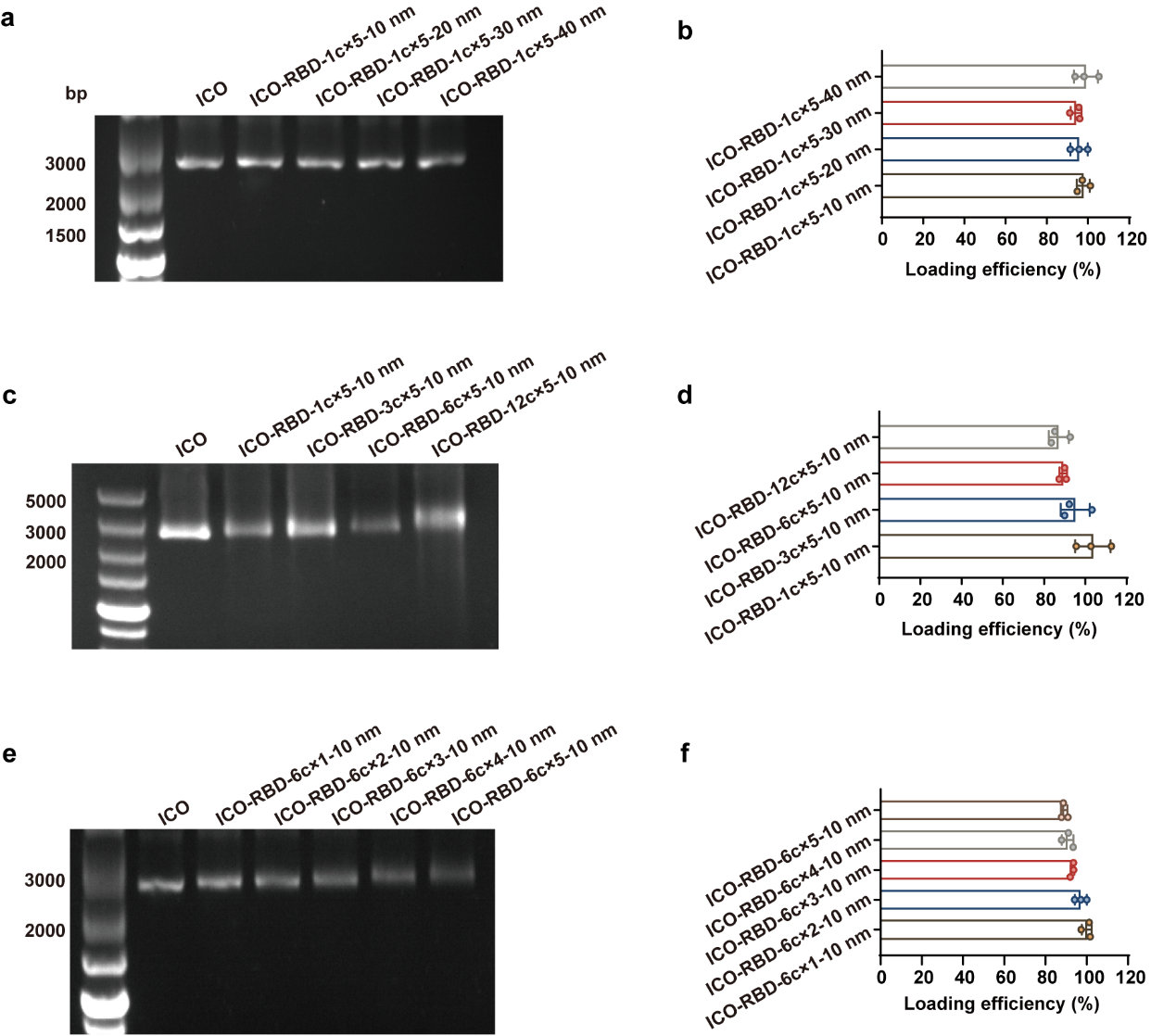

**Supplementary Figure 3.** Agarose gel electrophoresis and percentages of antigen coverage of ICO-RBD nanovaccines with the indicated surface antigen patterns. **(a-b)** Electrophoresis characterization (a) and percentages of antigen coverage (b) of the ICO-RBD nanovaccines with the indicated RBD spacing (n = 3). **(c-d)** Electrophoresis characterization (c) and loading efficiency (d) of the ICO-RBD nanovaccines with 1–12 RBD clusters of 5 RBD copies within clusters and 10 nm RBD spacing (n = 3). **(e-f)** Electrophoresis characterization (e) and percentages of antigen coverage (f) of the ICO-RBD-6c×n-10 nm nanovaccines with 1–5 antigen copies within clusters (n = 3). The data were processed on GraphPad Prism 8 and are presented as the mean ± SD.

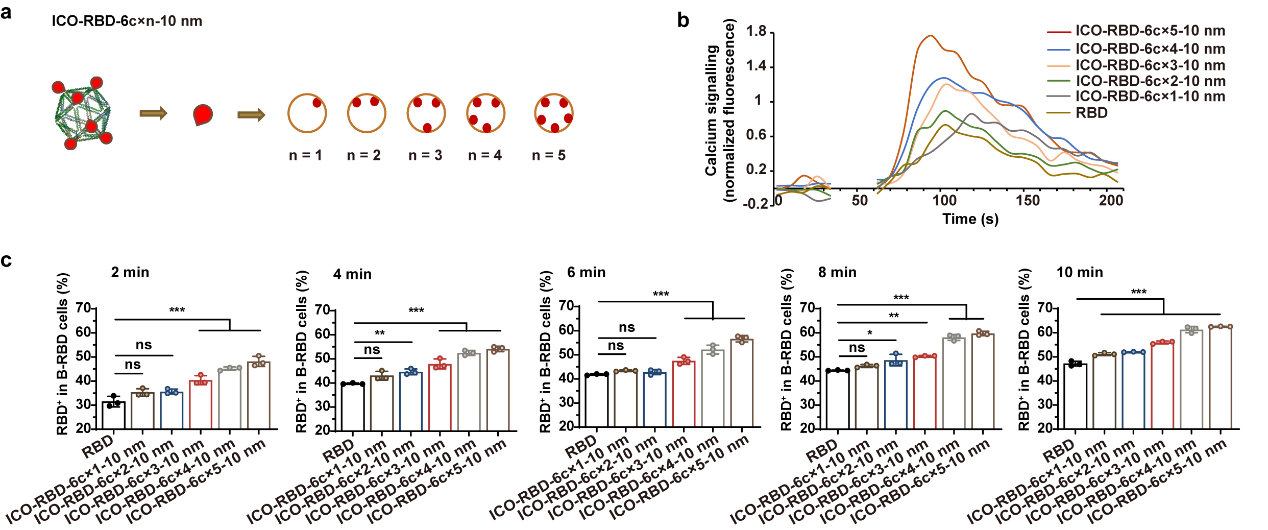

**Supplementary Figure 4.** Effect of the number of antigen copies within clusters on the ability of ICO-RBD nanovaccines to activate B-RBD cells. **(a)** Design of ICO-RBD nanovaccines with 6 RBD clusters of 1–5 RBD copies within clusters and 10 nm RBD spacing. **(b)** Ca^2+^ traces in B-RBD cells triggered by SpyCatcher-RBD proteins (RBD) or ICO-RBD nanovaccines with the indicated antigen copies within the clusters, as detected by Fluo-4 AM and flow cytometry. **(c)** Binding affinity between SpyCatcher-RBD proteins (RBD) or ICO-RBD nanovaccines with the indicated number of antigen copies within clusters and B-RBD cells (n = 3). The SpyCatcher-RBD proteins (RBD) or ICO-RBD nanovaccines bound to B-RBD cells were detected using anti-His antibodies and flow cytometry. The data were processed on GraphPad Prism 8 and are presented as the mean ± SD. Statistical significance (*P* value) was calculated by one-way ANOVA followed by Tukey’s test. ^*^, *P* < 0.05; ^**^, *P* < 0.01; ^***^, *P* < 0.001. ns, *P* > 0.05, no significant difference.

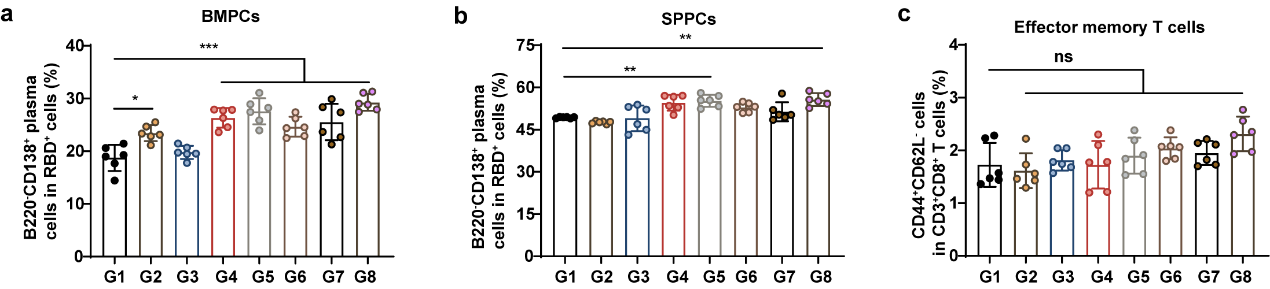

**Supplementary Figure 5.** Immune cell analysis at week 14 (cf. Figure 3). **(a-b)** Percentages of BMPCs (B220^-^CD138^+^) in RBD-specific bone marrow cells (a) and SPPCs (B220^-^CD138^+^) in RBD-specific splenocytes (b) at week 14, as detected using flow cytometry (n = 6). **(c)** Percentages of effector memory T cells (CD44^+^CD62L^-^) in CD3^+^CD8^+^ T cells in splenocytes at week 14, as detected using flow cytometry (n = 6). The data were processed on GraphPad Prism 8 and are presented as the mean ± SD. Statistical significance (*P* value) was calculated by one-way ANOVA followed by Tukey’s test. ^*^, *P* < 0.05; ^**^, *P* < 0.01; ^***^, *P* < 0.001. ns, *P* > 0.05, no significant difference.

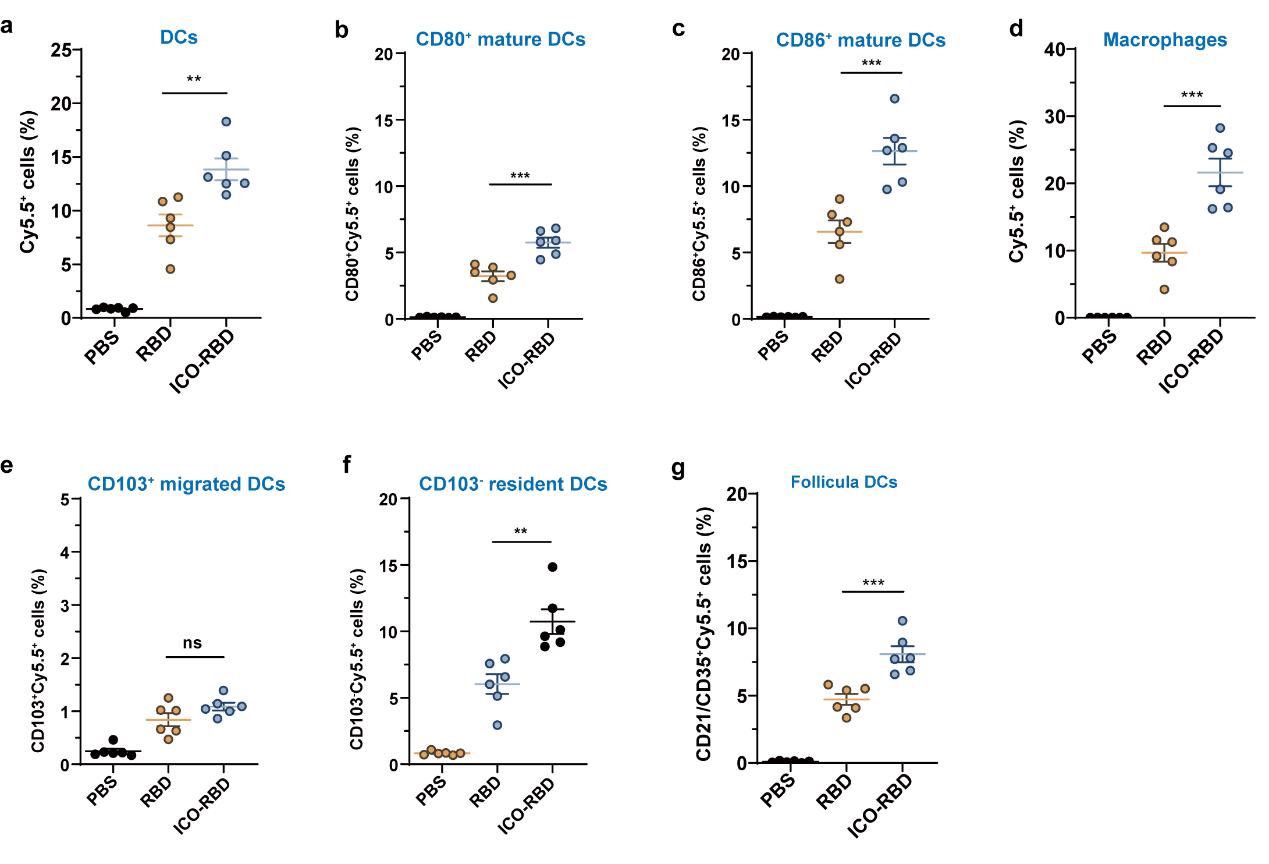
**Supplementary Figure 6.** Proportion of Cy5.5^+^ cells in different cell populations within inguinal lymph nodes at 12 h after immunization of Cy5.5-labeled RBD monomers or ICO-RBD nanovaccines, as detected using flow cytometry. **(a)** Percentages of Cy5.5^+^ DCs in CD11C^+^ DCs (n = 6). **(b)** Percentages of CD80^+^Cy5.5^+^ mature DCs in CD11C^+^ DCs (n = 6). **(c)** Percentages of CD86^+^Cy5.5^+^ mature DCs in CD11C^+^ DCs (n = 6). **(d)** Percentages of Cy5.5^+^ macrophages in F4/80^+^ macrophages (n = 6). **(e)** Percentages of CD103^+^Cy5.5^+^ migrated DCs in CD11C^+^ DCs (n = 6). **(f)** Percentages of CD103^-^Cy5.5^+^ resident DCs in CD11C^+^ DCs (n = 6). **(g)** Percentages of CD21/CD35^+^Cy5.5^+^ follicular DCs in CD11C^+^ DCs (n = 6). The data were processed on GraphPad Prism 8 and are presented as the mean ± SD. Statistical significance (*P* value) was calculated by one-way ANOVA followed by Tukey’s test. ^**^, *P* < 0.01; ^***^, *P* < 0.001. ns, *P* > 0.05, no significant difference.

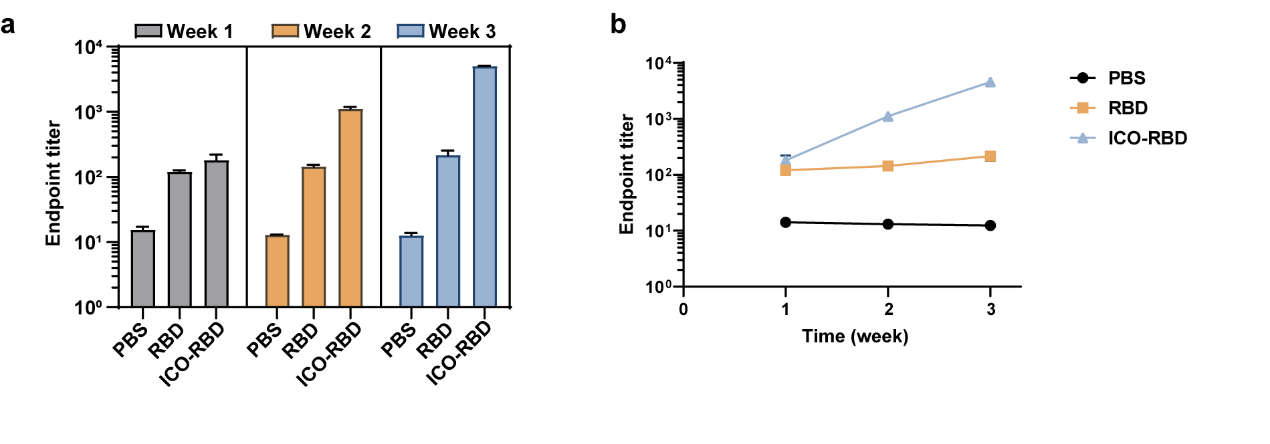

**Supplementary Figure 7.** SARS-CoV-2 RBD-specific IgG titers in serum after priming vaccination. Mice were randomly divided into 3 groups and serum was collected every week. **(a)** RBD-specific IgG titers (n = 8). **(b)** Time-titer curves (n = 8). The data were processed on GraphPad Prism 8 and are presented as the mean ± SEM.

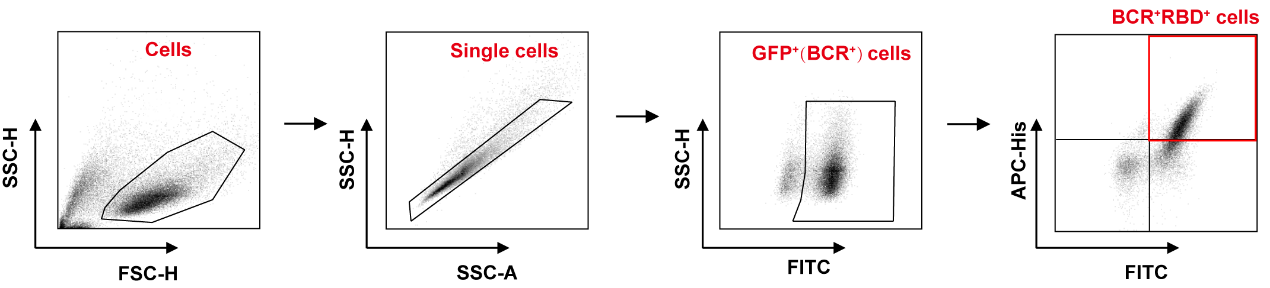

**Supplementary Figure 8.** Gating strategy for RBD^+^ cells in B-RBD cells in Figure 2b, Figure 2e and Supplementary Figure 4c.

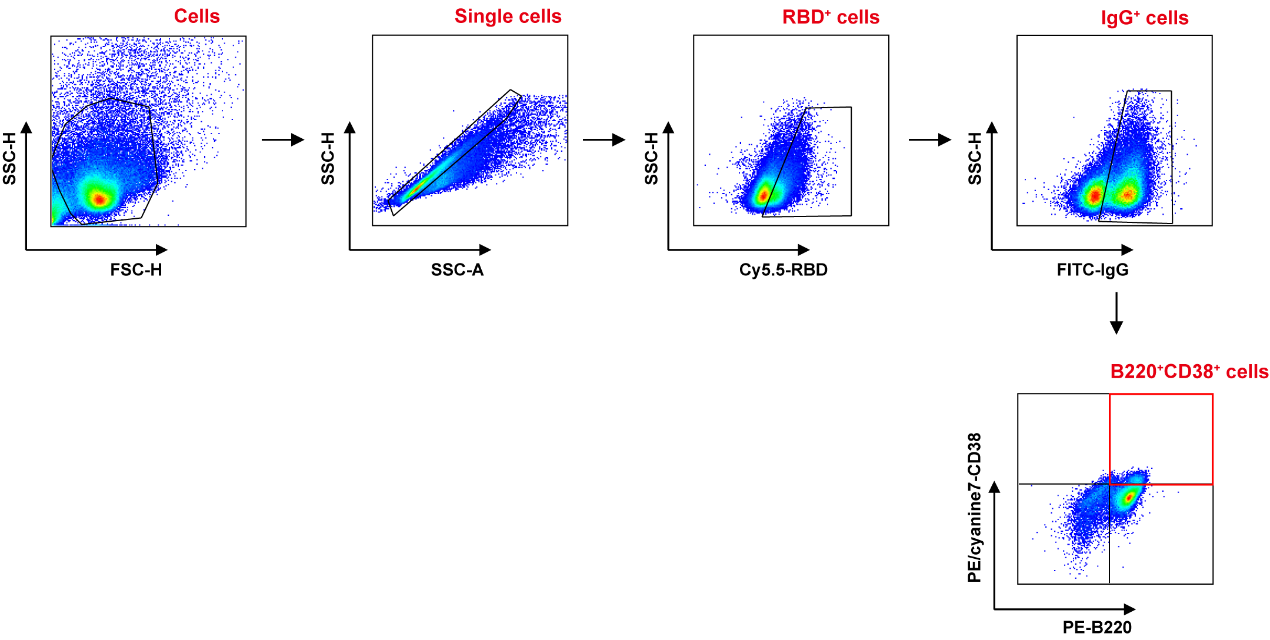

**Supplementary Figure 9.** Gating strategy for MBCs (B220^+^CD38^+^) in RBD^+^IgG^+^ cells in splenocytes in Figure 3g.

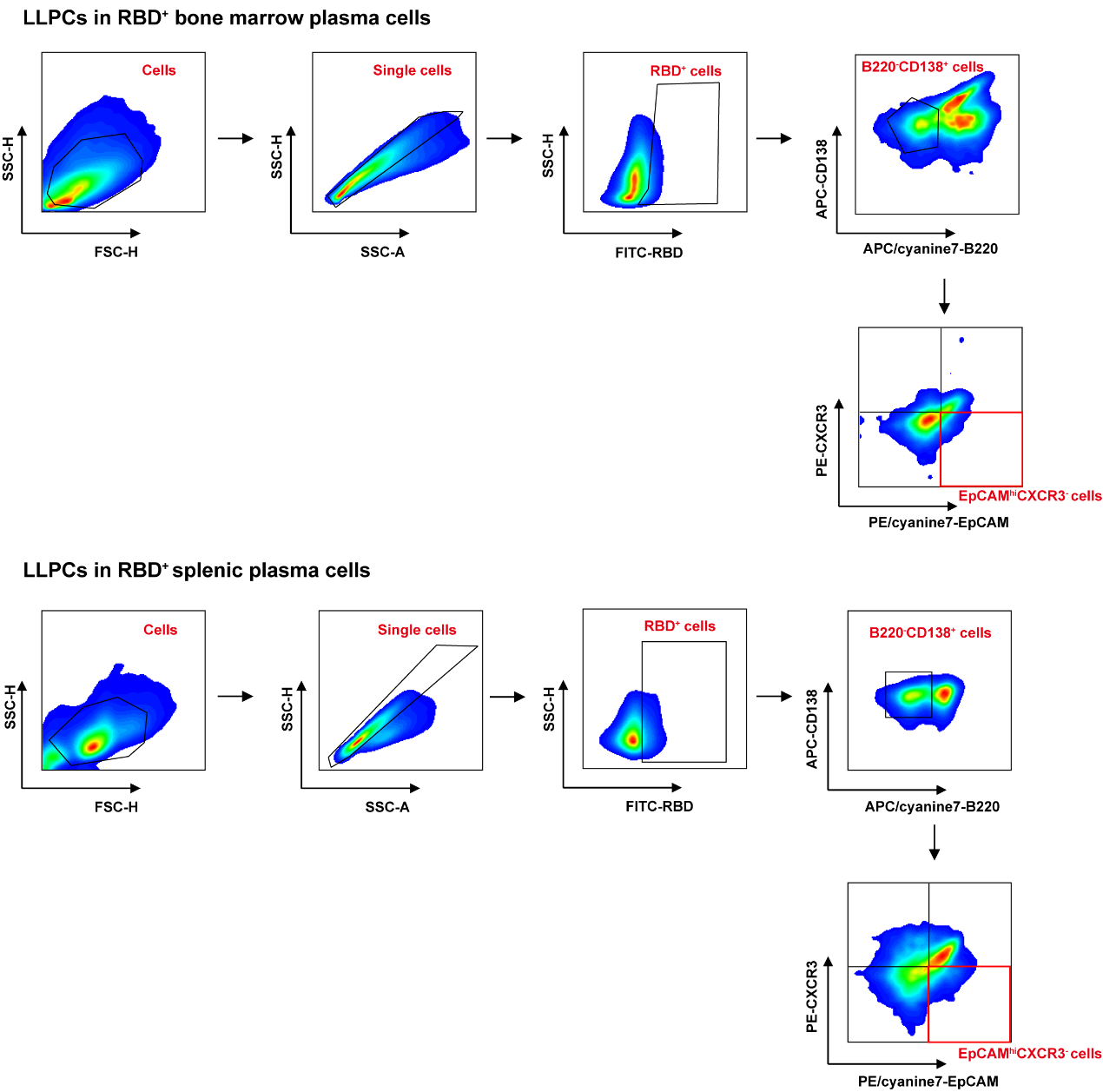

**Supplementary Figure 10.** Gating strategy for LLPCs (EpCAM^hi^CXCR3^−^) in RBD^+^ bone marrow plasma cells (BMPCs, B220^-^CD138^+^) and RBD^+^ splenic plasma cells (SPPCs, B220^-^CD138^+^) in Figures 3h-i.

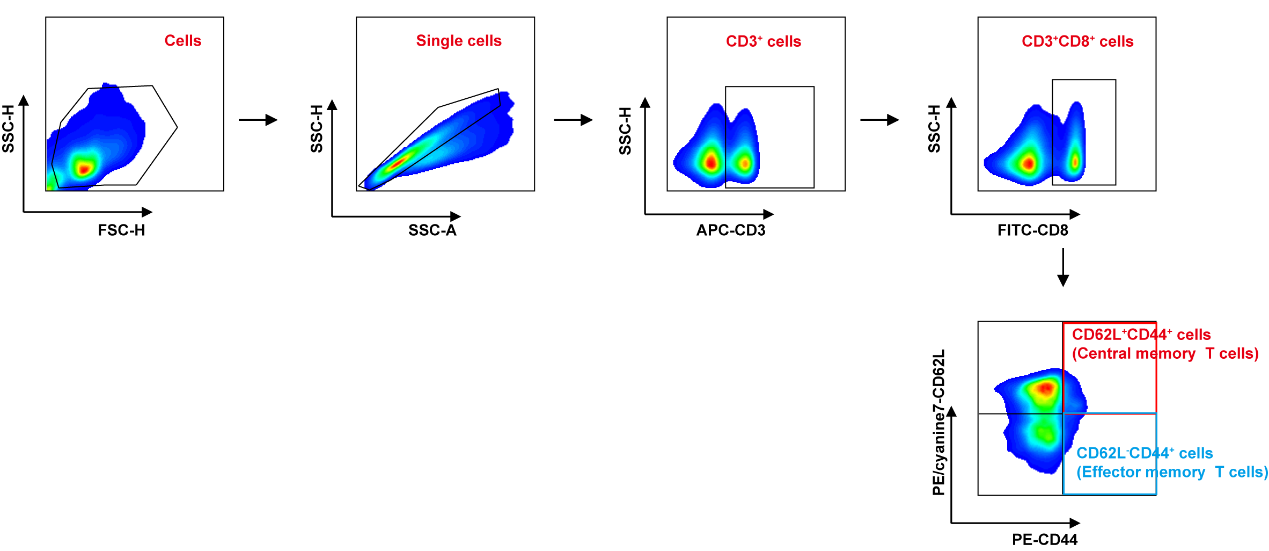

**Supplementary Figure 11.** Gating strategy for central memory T cells (CD44^+^CD62L^+^) in CD3^+^CD8^+^ T cells in Figure 3j and effector memory T cells (CD44^+^CD62L^-^) in CD3^+^CD8^+^ T cells in Supplementary Figure 5c.

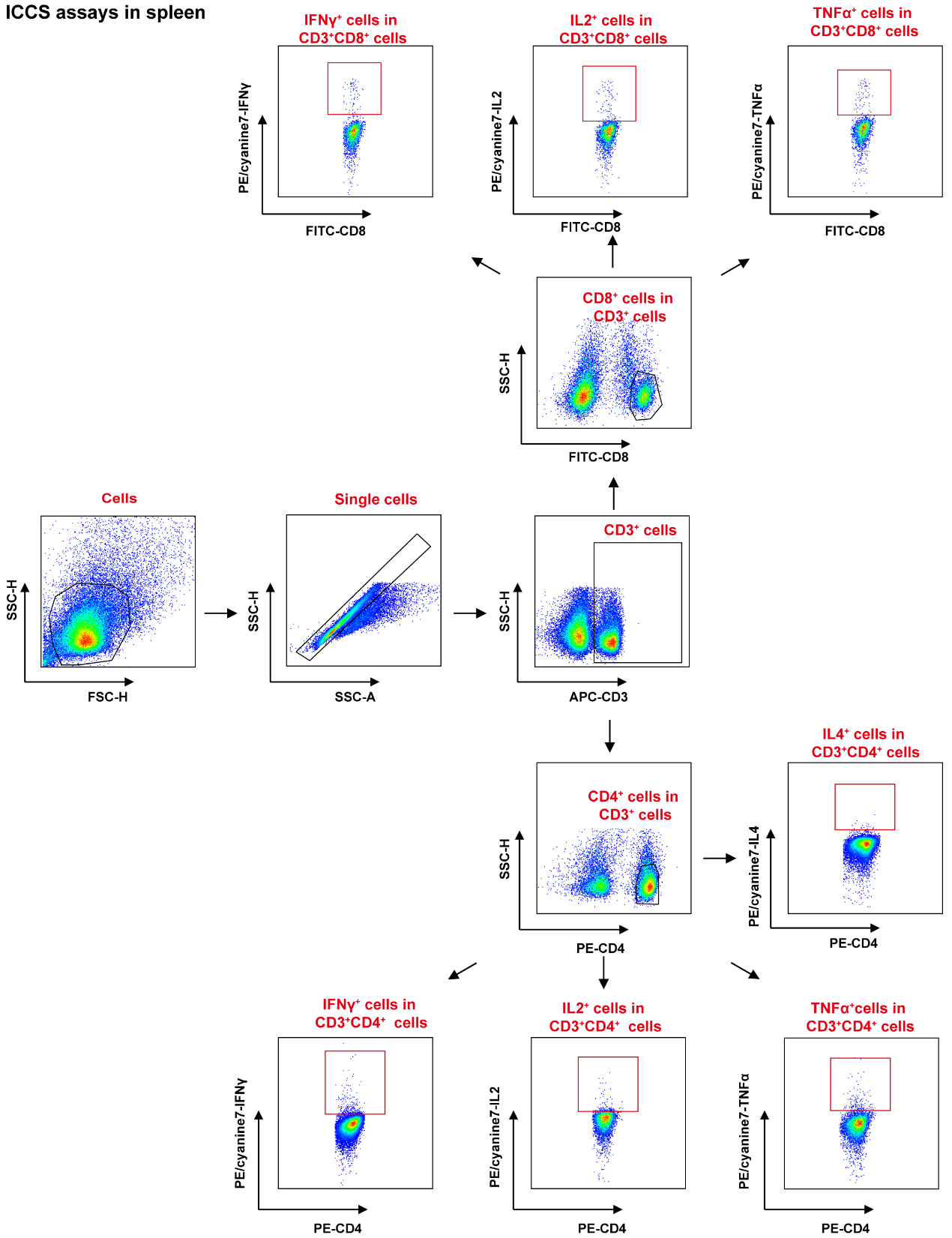

**Supplementary Figure 12.** Gating strategy for IFN-γ^+^/IL-2^+^/TNF-α^+^ cells in CD3^+^CD8^+^ T cells and CD3^+^CD4^+^ T cells in the spleen by intracellular cytokine staining (ICCS) assays in Figures 4a-b and IL-4^+^ cells in CD3^+^CD4^+^ T cells within the spleen in Figure 4e.

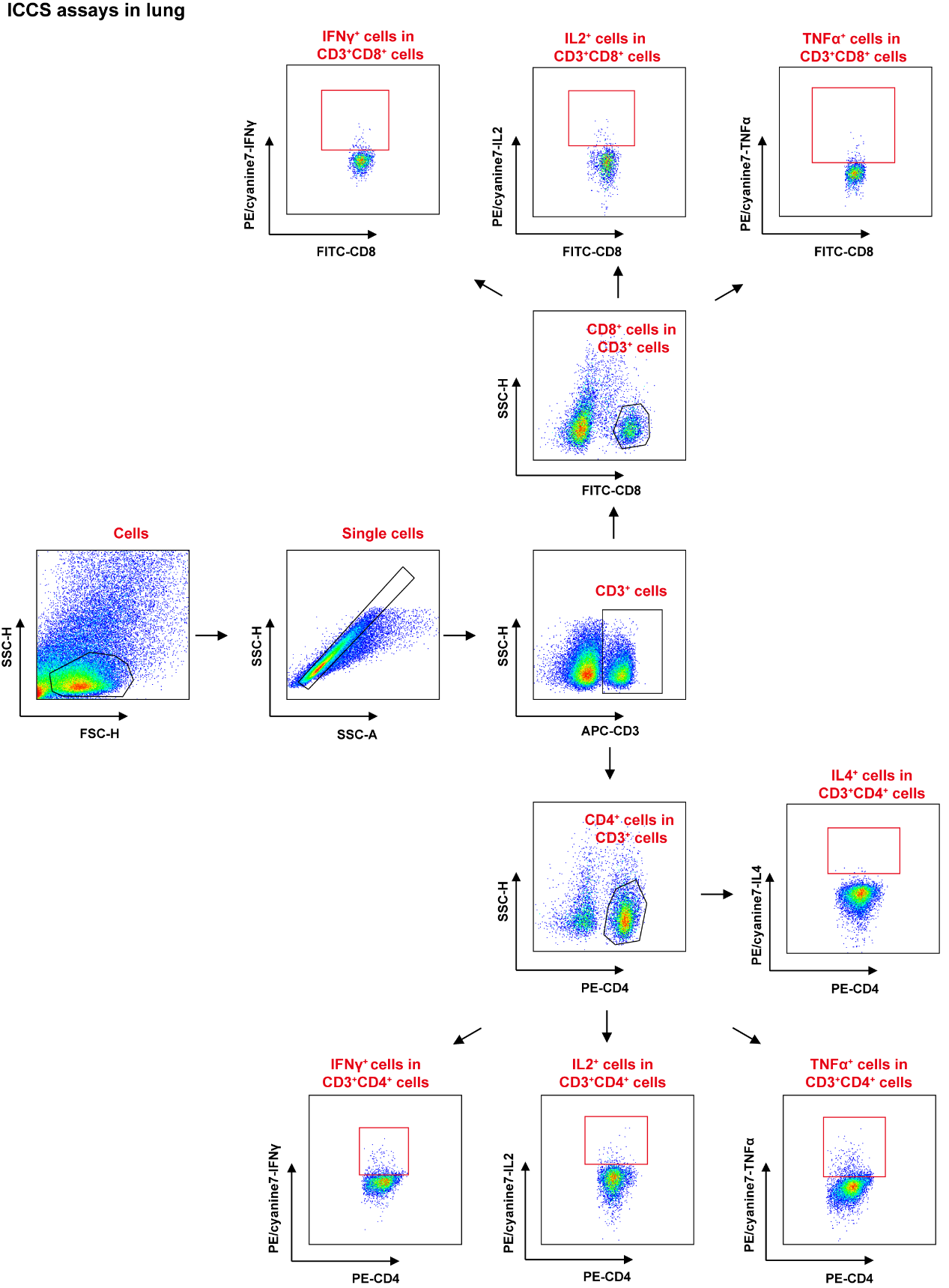

**Supplementary Figure 13.** Gating strategy for IFN-γ^+^/IL-2^+^/TNF-α^+^ cells in CD3^+^CD8^+^ T cells and CD3^+^CD4^+^ T cells within the lung by ICCS assays in Figures 4c-d and IL-4^+^ cells in CD3^+^CD4^+^ T cells within the lung in Figure 4f.

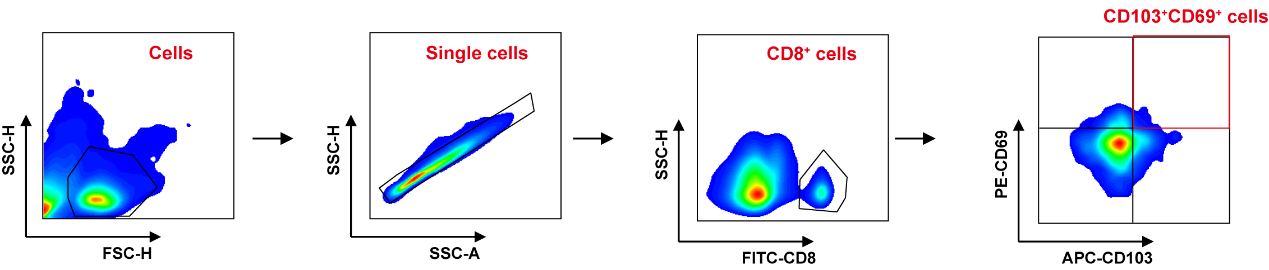

**Supplementary Figure 14.** Gating strategy for CD69^+^CD103^+^ TRMs in CD8^+^ T cells in the lung in Figure 4g.

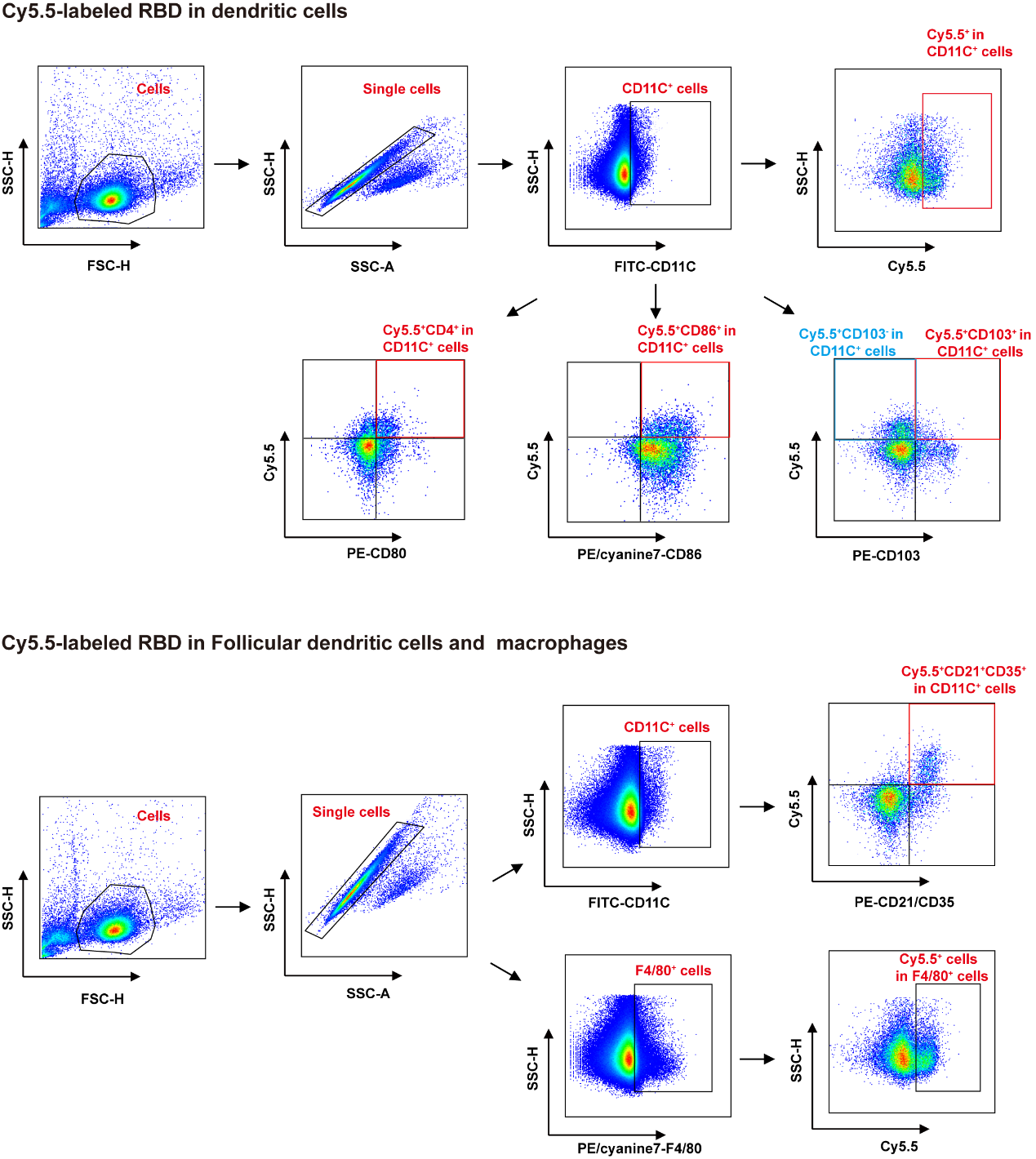

**Supplementary Figure 15.** Gating strategy for Cy5.5^+^ DCs, CD80/CD86^+^Cy5.5^+^ matured DCs, CD103^+^Cy5.5^+^ migrated DCs and CD103^-^Cy5.5^+^ resident DCs in CD11C^+^ DCs, CD21/CD35^+^Cy5.5^+^ follicular DCs in CD11C^+^ DCs, Cy5.5^+^ macrophages in F4/80^+^ macrophages in the inguinal lymph nodes in Supplementary Figure 6.

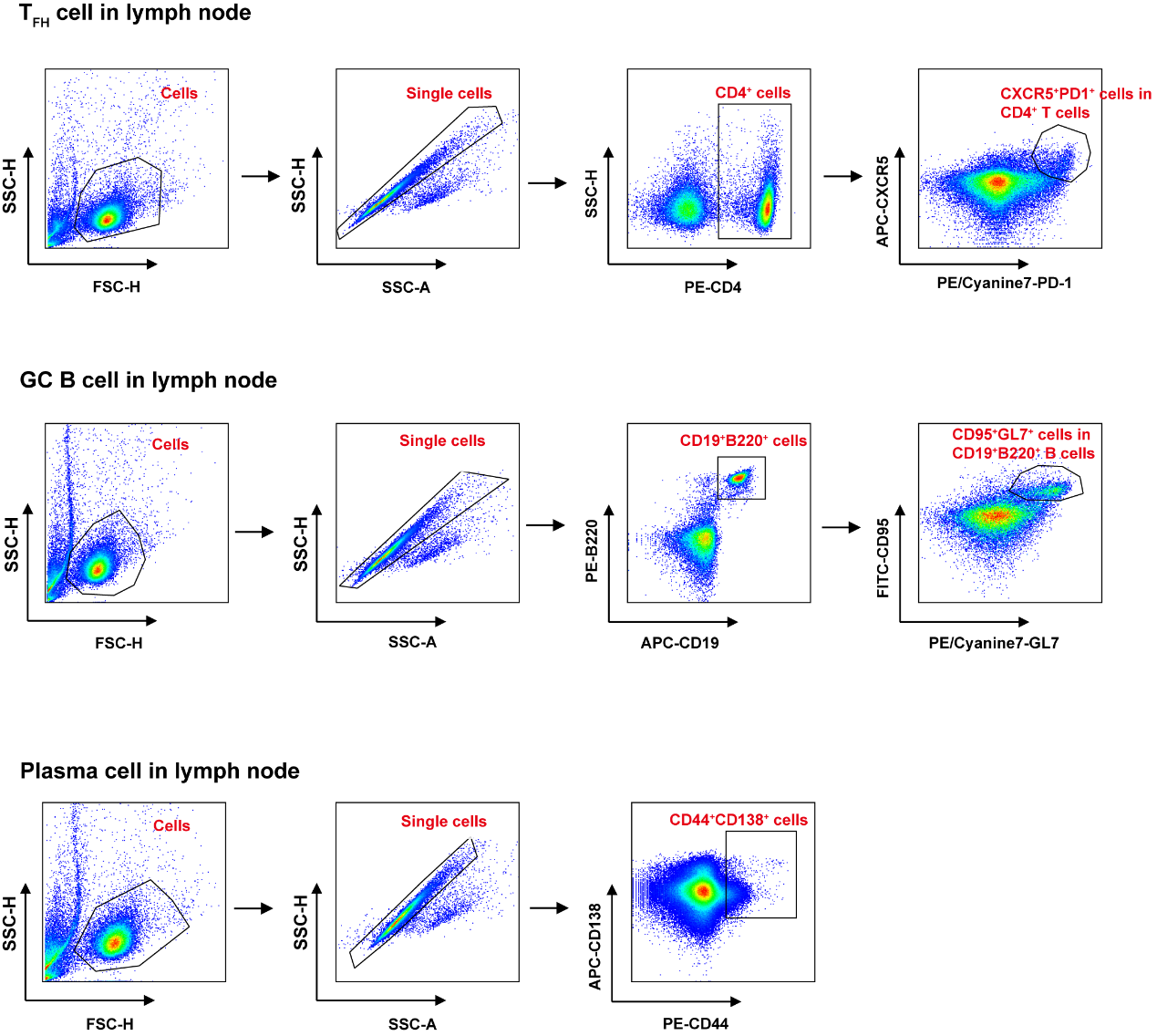

**Supplementary Figure 16.** Gating strategies for T_FH_ cells (CXCR5^+^PD1^+^) in CD4^+^ cells, GC B cells (CD95^+^GL7^+^) in CD19^+^B220^+^ cells and plasma cells (CD44^+^CD138^+^) in Figure 5e.
